# An mRNA-encoded, long-lasting Interleukin-2 restores CD8^+^ T cell neoantigen immunity in MHC class I-deficient cancers

**DOI:** 10.1101/2023.07.18.549445

**Authors:** Jan D. Beck, Mustafa Diken, Martin Suchan, Michael Streuber, Elif Diken, Laura Kolb, Lisa Allnoch, Fulvia Vascotto, Daniel Peters, Tim Beißert, Özlem Türeci, Sebastian Kreiter, Mathias Vormehr, Ugur Sahin

## Abstract

MHC class I antigen presentation deficiency is considered to be the most prevalent cancer immune escape mechanism. Despite its increasing occurrence, the mechanistic implications, and potential strategies to address this challenge, remain poorly understood. Studying β2-microglobulin (B2M) deficient mouse tumor models, we found that MHC class I loss leads to a substantial immune desertification of the tumor microenvironment (TME) and broad therapeutic resistance to immune-, chemo- and radiotherapy. We show that treatment with long-lasting mRNA-encoded interleukin-2 (IL2) restores an immune cell infiltrated, IFNγ-promoted, highly proinflammatory TME signa-ture, and when combined with a tumor-targeting monoclonal antibody (mAb), can overcome ther-apeutic resistance. Surprisingly, we identified that effectiveness of this treatment is driven by ne-oantigen-specific IFNγ-releasing CD8^+^ T cells that recognize neoantigens cross-presented by TME-resident activated macrophages that under IL2 treatment acquire augmented antigen presen-tation proficiency along with other M1-phenotype-associated features. Our findings highlight the unexpected importance of restoring neoantigen-specific immune responses in the treatment of cancers with MHC class I deficiencies.

## Introduction

Loss of antigen presentation through MHC class I deficiency is a key mechanism of cancer immune escape from immune surveillance and killing by cytotoxic CD8^+^ T lymphocytes. Various molecular mechanisms contribute to MHC class I deficiency, including somatic mutations and epigenetic modifications in genes involved in antigen presentation, or tumor microenvironmental factors such as hypoxia (1). Throughout the life cycle of a tumor, immunosurveillance acts as a selective pressure towards the emergence of MHC class I-deficient cancer cells (2,3). Cancer immunotherapy, particularly the broad use of immune checkpoint inhibitors, such as PD-1 or CTLA- 4 blocking antibodies, is associated with an increased emergence of MHC class I deficient tumors with either inactivating mutations in MHC class I molecules, in genes involved in interferon (IFN) response pathways (4–6), or in *B2M* encoding β2-microglobulin (B2M) (6–12). Understanding the consequences of MHC class I deficiency and development of therapeutic strategies to circumvent and address these challenges is essential for improving the outcomes of cancer immunotherapy.

### Immune desertification and cross-modality therapy resistance of MHC class I-deficient tumors

B2M is the shared component of all MHC class I molecules. Its loss results in complete absence of MHC class I surface expression and consequently the lack of CD8^+^ T-cell recognition. To study MHC class I presentation deficiency in different settings we deleted *B2m* genes in three mouse tumor cell lines. Of these, CT26 and MC38 form highly immunogenic tumors and elicit spontaneous CD8^+^ T-cell responses (13,14), whereas B16F10 melanomas have a lower prevalence of tumor-infiltrating leukocytes (TIL) and are considered non-immunogenic tumors (13). *B2m*-knock-out (*B2m^-/-^*) cells lack MHC class I surface expression and therefore are not recognized by co-cultured antigen-specific CD8^+^ T cells (Fig. S1, A and B). Analyzing immune cell infiltration longitudinally during growth of wild-type and *B2m^-/-^* tumors in syngeneic mice, we observed a gradual decrease in immune cell infiltrates in CT26-*B2m^-/-^* tumors and immune cell desertification of the tumor microenvironment (TME) within 20 days after inoculation (Fig. 1A) with decreased CD8^+^ T cells, NK cells and conventional type I dendritic cells (cDC1) (Fig. S1C). B2M loss had similar effects on MC38 tumors, while B16F10 tumors, as expected, regardless of their *B2m* genotype exhibited sparse immune infiltration (Fig. S1, C and D).

**Figure 1.**
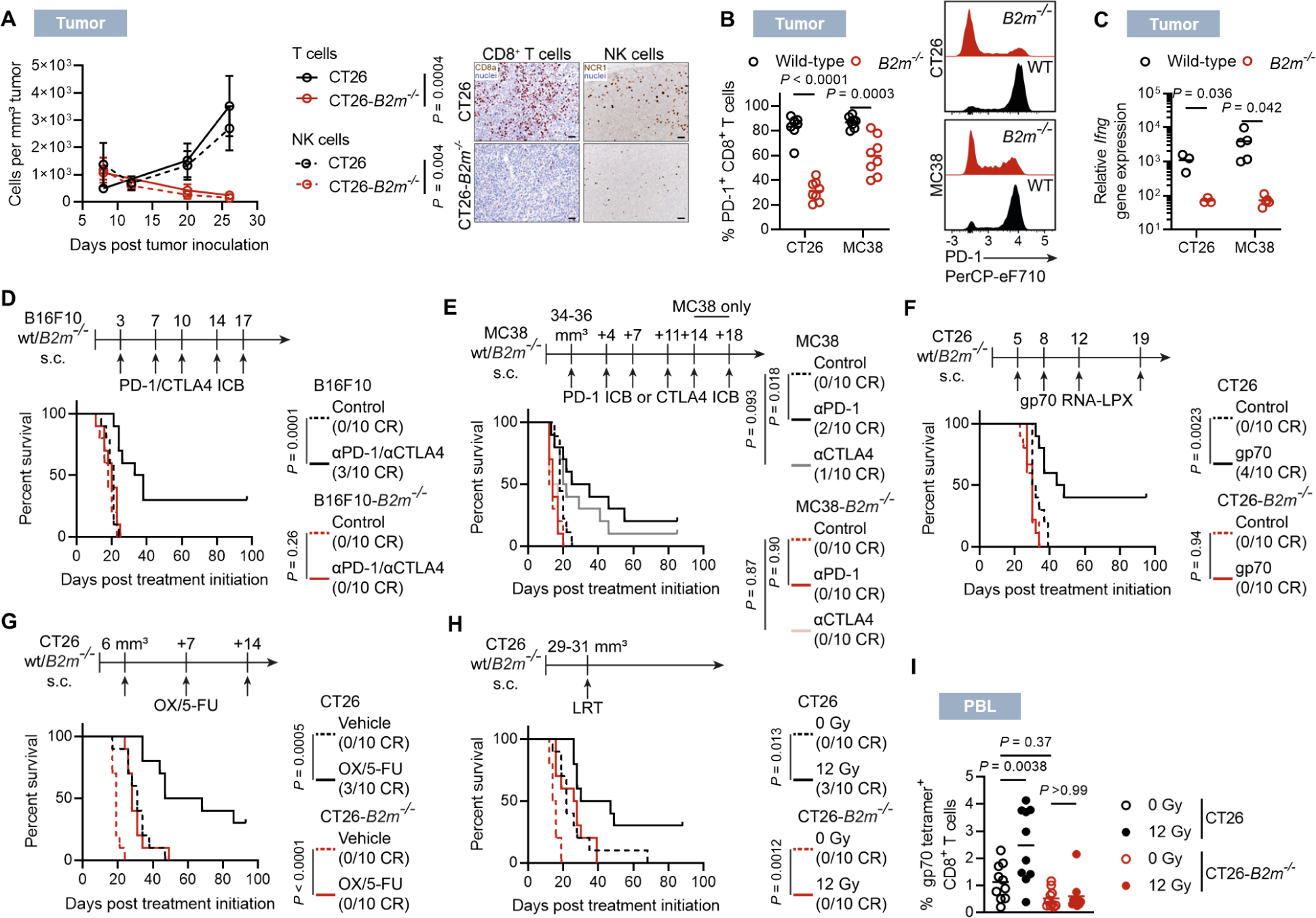
Immune desertification and therapy resistance of MHC class I-deficient tumors. (**A**) Longitudinal dynamics of T- and NK-cell infiltration in CT26 or CT26-*B2m^-/-^* tumors and immunohistochemical (IHC) analysis of T- and NK-cell infiltration 19 days after inoculation. Scale bar = 50 µm. (**B**) PD-1 expression in CT26 or MC38 wild-type or *B2m^-/-^*-infiltrating CD8^+^ T cells 20 days after inoculation. (**C**) *Ifng* expression in CT26 or MC38 wild-type or *B2m^-/-^* tumor tissues 19 days after inoculation. (**D** to **H**) Survival of mice bearing the indicated parental or *B2m^-/-^* tumor variants upon treatment with αPD-1/αCTLA4 ICB combination or isotype controls (D), αPD-1, αCTLA4 or an irrelevant control mAb (E), gp70-encoding mRNA-LPX vaccine or empty-vector mRNA-LPX as control (F), Oxaliplatin/5-fluorouracil (OX/5-FU) doublet or vehicle control (G), local radiotherapy (LRT) at a dose of 12 Gy or 0 Gy as control (H). (**I**) gp70 antigen-specific CD8^+^ T cells in the blood 9 d after LRT (H). n=4-5 per time point (A; left) and representative IHC stainings (A; right). n=8 (B). n=3 (CT26) and n=5 (MC38) (C). n=10 (D-I)

The changes in the immune cell composition were limited to the TME and not seen in tumor-draining lymph nodes (TDLNs), with the exception of a minor decrease in the quantity of cDC1 residing in the TDLNs of mice carrying MC38-*B2m*^-/-^ tumors (Fig. S1E).

CT26-*B2m^-/-^* and MC38-*B2m^-/-^* tumors showed significantly faster progression *in vivo* but not under cell culture conditions *in vitro* (Fig. S1, F and G), while B16F10 tumors did not show significant growth differences compared to their wild-type counterparts (Fig. S1F). The fraction of PD- 1-expressing CD8^+^ T cells was significantly reduced by B2M loss in both models (Fig. 1B) and mice with CT26-*B2m^-/-^* tumors had a significantly lower frequency of CD8^+^ T cells specific for the immunodominant endogenous retroviral antigen gp70 (15) (Fig. S2A).

In CT26-*B2m^-/-^* and MC38-*B2m^-/-^* tumors, expression of *Il2*, which promotes proliferation of immune cells, and *Ifng*, an effector molecule released by CD8^+^ T cells upon cognate antigen recognition, were significantly reduced (Fig. 1C and fig. S2B). In line with this, *B2m^-/-^* tumors showed reduced levels of the IFNγ-regulated, T and NK cell attracting chemokines *Cxcl9* and *Cxcl10* (16,17). The expression of *Xcl1*, which is produced by NK cells and contributes to the recruitment of cDC1, was also diminished (Fig. S2B) (18). Consistent with this reduced inflammatory signature in the TME, PD-L1 was downregulated in CT26-*B2m^-/-^* and MC38-*B2m^-/-^* tumor cells *in vivo* (Fig. S2C), but not in cultured MC38-*B2m^-/-^* and CT26-*B2m^-/-^* cells (Fig. S2D). The IFNγ-inducible markers PD-L1 (19) and MHC class II (20) were significantly downregulated in CT26-*B2m^-/-^* and MC38-*B2m^-/-^* tumor-infiltrating macrophages (Fig. S2E), indicating a shift from an inflammatory M1-like phenotype to a more immunosuppressive phenotype.

Having revealed the profound changes in TME contexture and accelerated *in vivo* growth of MHC class I-deficient tumors that cannot be directly recognized by CD8^+^ T cells, we investigated the impact of B2M deficiency on the effects of different treatments. Mutation of *B2M* has been associated with resistance of human cancers to T-cell based therapies such as immune checkpoint blockade (ICB) and mRNA-based neoantigen vaccines. In line with those observations in patients, treatment of B16F10 and of MC38 tumor-bearing mice with PD-1 or CTLA4 blocking antibodies resulted in tumor rejection and significantly prolonged survival, but completely lacked activity against the respective *B2m^-/-^* variants (Fig. 1, D and E and fig. S3, A and B). Treatment of CT26- bearing mice with an mRNA-lipoplex (RNA-LPX) vaccine encoding the immunodominant antigen gp70, which was demonstrated previously to induce strong expansion of antigen-specific cytotoxic CD8^+^ T cells (21,22), prolonged survival and induced rejection of 40% of the CT26 tumors, whereas CT26-*B2m^-/-^* tumors progressed unaffectedly (Fig. 1F and fig. S3C).

Furthermore, we assessed the responsiveness of MHC class I-deficient tumors to chemotherapy and local radiotherapy (LRT) that are considered to exert their anti-tumor activity primarily by direct cytotoxicity. Treatment with a platinum-based chemotherapy improved the survival of mice bearing MC38 or CT26 tumors, but less so of mice bearing the *B2m^-/-^*versions of these tumors (Fig. 1G and fig. S4, A to E). Only the wild-type tumors were rejected, and in the *B2m^-/-^* variants growth reduction was less profound. Similarly, the survival-prolonging effect of LRT was more pronounced in mice with CT26 than in those with CT26-*B2m^-/-^* tumors (Fig. 1H and fig. S4, F and G). Tumor rejection was observed exclusively in mice bearing wild-type tumors, whereas the impact of LRT on growth of *B2m^-/-^* tumors was modest and all tumors eventually progressed. CD8^+^ T cells recognizing the immunodominant gp70 antigen were expanded in the blood of locally irradiated mice bearing wild-type but not *B2m^-/-^* CT26 tumors (Fig. 1I).

Notably, even tumor antigen-targeted monoclonal antibodies (mAb)s, utilizing MHC class I independent cognate mechanisms for tumor eradication, exhibited only a marginal therapeutic efficacy (shown below).

Our findings indicate that loss of MHC class I antigen presentation and the resulting immune desertification of the TME promote resistance to diverse therapeutic interventions.

### Reversion of resistance of MHC class I-deficient tumors to various treatment modalities by mAb/IL2 therapy

Interleukin-2 (IL2) cytokine therapy can enhance T-cell infiltration into tumors by promoting the activation, expansion, and survival of immune cells (23). Recent studies have reported the treatment of MHC class I-deficient tumors using IL2 variants specifically designed to bind the low-affinity IL2-receptor complex (common γ-chain/IL2Rβ), thereby promoting NK-cell functionality (24,25). These reports, together with the IL2 downregulation that we observed in immune deserted *B2m^-/-^* tumors (Fig. S2B), motivated us to assess the impact of IL2 administration. Unlike some of the previous studies, we opted for wild-type IL2 to ensure optimal stimulation of T cells (26).

Recombinant IL2 has an unfavorable pharmacokinetic profile that is reflected by a short serum half-life. Frequent administration at high dose levels is required, which is associated with profound toxicity (27,28). We engineered a lipopolyplex-formulated mRNA-encoded IL2 as well as an IL2 fused to Albumin. As previously shown, intravenously administered lipopolyplexed mRNA is subject to highly specific delivery to the liver and expression in liver cells. Minor mRNA translation occurs in spleen and lymph nodes whereas no translation is observed in other organs, e.g. lungs (29). The mRNA-encoded protein product is released from liver cells into the blood stream (Fig. 2A). Compared to intravenously administered recombinant IL2 protein, IL2 translated *in vivo* from mRNA reached robust serum concentrations over an extended period of time in mice, and the mRNA-encoded Albumin-fused IL2 version had an even further prolonged serum half-life (Fig. 2A).

**Figure 2.**
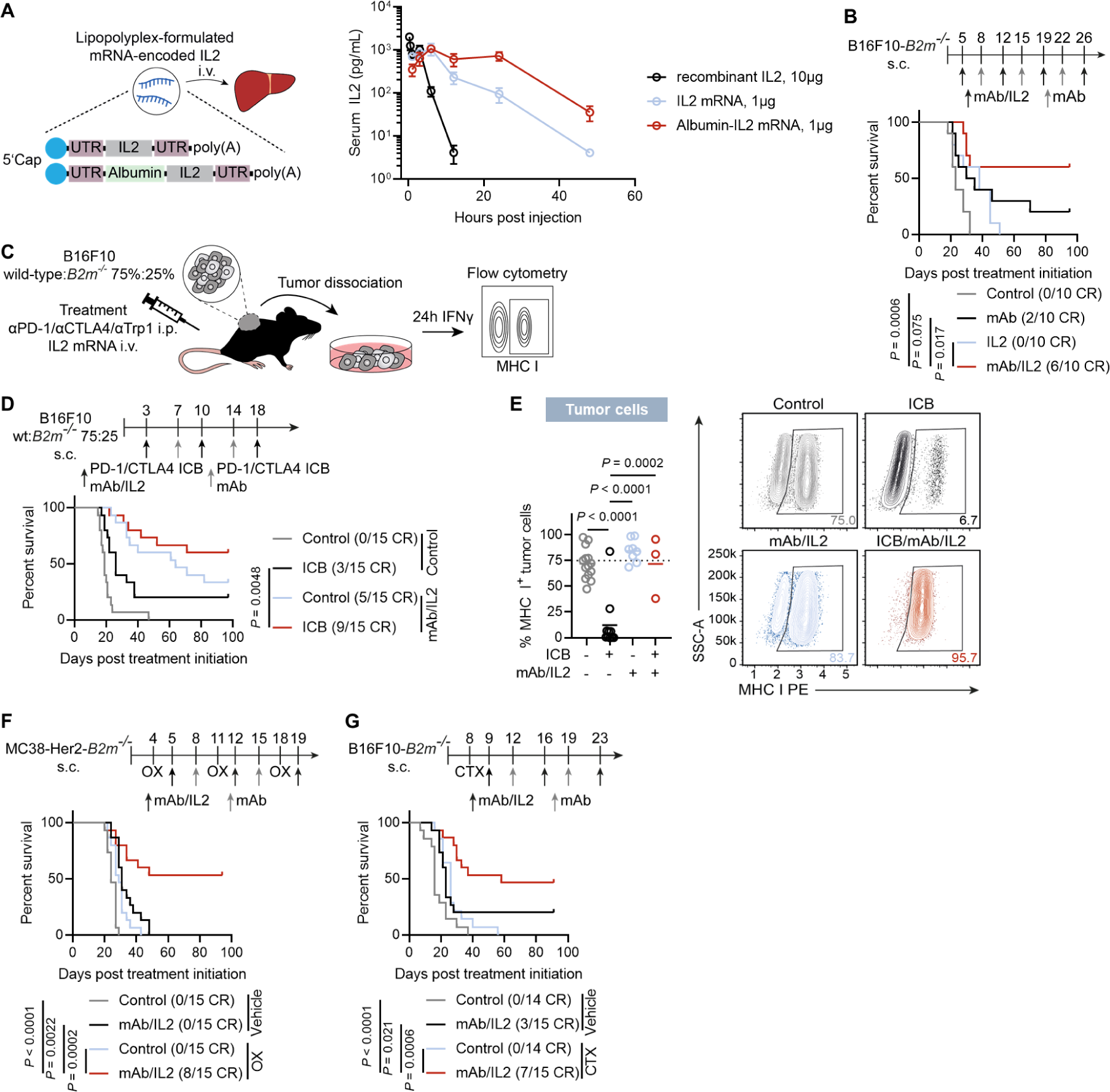
Reversion of resistance of MHC class I-deficient tumors to various treatment modalities by mAb/IL2 therapy. (**A**) Illustration of IL2 and Albumin-IL2 mRNA including 5’ Cap, 5’ and 3’ untranslated regions (UTR)s flanking the open reading frames and poly(A)-tail (left) and pharmacokinetics of mRNA-delivered long-acting IL2 compared to recombinant protein (right). (**B**) Survival of mice treated with αTrp1 mAb and intravenous IL2 mRNA-lipopolyplex (short IL2) or either of the compounds. (**C** to **E**) Mice were inoculated with B16F10 wild-type cells spiked with B16F10-*B2m*^-/-^ cells (75%:25%) and treated with αTrp1 mAb/IL2, PD-1 and CTLA4 combination ICB or the quadruple combination. Tumors were collected when mice reached endpoint criteria and the proportion of MHC class I^+^ tumor cells was determined after 24 h stimulation with IFNγ. Experimental scheme (C), survival of tumor-bearing mice (D) and quantification of MHC class I^+^ tumor cells (Dashed line represents the fraction of MHC class I^+^ cells at the time of inoculation) (E). (**F** and **G**) Survival of mice bearing MC38-Her2-*B2m*^-/-^ tumors after treatment with αHer2 mAb/IL2, oxaliplatin (OX) or the triple combination (F) or B16F10-*B2m*^-/-^ tumors after treatment with αTrp1 mAb/IL2, cyclophosphamide (CTX), the triple combination or controls (G). Albumin-encoding mRNA and isotype mAbs served as controls across all experiments. n=3 (A). n=10 (B). n=15 (D). n=3 (ICB/mAb/IL2), n=11 (ICB), n=8 (mAb/IL2) and n=14 (control) (E). n=15 (F). n=14-15 (G).

Therapeutic doses of the mRNA-encoded Albumin-fused IL2, as compared to mRNA-encoded Albumin alone, resulted in the expected T-cell activation-related serum cytokine signature (upregulation of e.g. IFNγ, TNFα), with rare modest increase of commonly adverse event-related inflammatory cytokines IL-1β and IL-6 (Fig. S5A). We performed histopathological assessment of livers (being the organ in which IL2 mRNA is translated), and lungs (a major organ of vascular leak syndrome manifestation as side effect of high-dose IL2 treatment). The livers of mRNA-encoded Albumin treated control mice displayed occasional sparse lobular (Fig. S5B), perivascular and periportal (Fig. S5C) immune cell infiltrates and single necrotic cells, that are background lesions commonly observed in mice (30). These were slightly more frequent in mice treated with mRNA-encoded Albumin-fused IL2. Manifest signs of inflammation or necrosis, elevation of transaminase above the reference range (Fig. S5D), increase of dehydrogenases (Fig. S5E) or clinical signs of liver malfunction did not occur. Lungs of mice from both treatment groups exhibited occasional perivascular immune cell infiltrates without signs of vascular damage that were slightly more pronounced in Albumin-IL2 mRNA treated mice (Fig. S5F) and lung weight increases (as proxy for vascular leak syndrome) were statistically insignificant (Fig. S5G). Mice treated with Albumin-IL2 mRNA showed no systemic clinical signs of treatment-related side effects and exhibited normal body weight gain (Fig. S6H).

Given its favorable pharmacokinetic profile, the long-acting Albumin-fused IL2 mRNA was used for all experiments presented below (referred to as ‘IL2’ hereafter for simplicity) and is currently being evaluated in an ongoing first-in-human phase I trial (alias BNT153; NCT04710043).

We used two mAb-targetable MHC class I-deficient tumor models to test IL2 in conjunction with tumor-specific mAb treatment, which we chose as an alternative way to mediate direct recognition of tumor cells lacking MHC class I by immune cells. One model involved MC38-*B2m^-/-^* cells stably expressing rat Her2/neu (MC38-Her2-*B2m^-/-^*, Fig. S6A) to be treated with anti (α)Her2 mAb. The other model was the Trp1-positive B16F10-*B2m^-/-^* cell line targetable by αTrp1 mAb.

Treatment of B16F10-*B2m^-/-^*tumors with mAb/IL2 combination therapy (3-4 doses Q1W) led to tumor rejection and long-term survival in 60% of mice and was significantly superior to either IL2 alone or mAb alone (Fig. 2B and fig. S6B). The treatment effect and superiority of the mAb/IL2 combination was even more pronounced in MC38-Her2-*B2m^-/-^*-bearing mice with a long-term survival of 50% as compared to 10% in IL2-only and 0% in mAb-only treated mice (Fig. S6, C and D). The mAb/IL2 combination was effective in MHC class I-positive tumors as well and showed robust antitumor activity in the wild-type B16F10 model (Fig. S6, E and F), with long-term survival of 30% of mice.

Re-challenge of mice cured from B16F10*-B2m^-/-^*tumors with B16F10*-B2m^-/-^* cells resulted in slower, yet progressive tumor growth, indicating that immunological memory established in the context of mAb/IL2 therapy is not fully protective (Fig. S6G).

Considering that human tumors are clonally heterogeneous and that in the clinical setting resistance is often acquired by outgrowth of subclonal MHC class I-deficient tumor cell populations, we inoculated mice with wild-type B16F10 cells that were spiked with B16F10-*B2m^-/-^* cells (wild-type:*B2m^-/-^* ratio of 75%:25%) (Fig. 2C). PD-1/CTLA4 ICB treatment of mice inoculated with these heterogeneous tumors had a poor effect on their survival. A 20% tumor rejection rate was observed, likely attributable to bystander elimination (31) of B16F10-*B2m^-/-^* cells (Fig. 2D and fig. S6H). In tumors of mice meeting endpoint criteria, the wild-type to *B2m^-/-^* tumor cell ratio remained unaltered in the non-active control group (Fig. 2E). In contrast, ICB-treated mice displayed an increase in the proportion of MHC class I-deficient cells from 25% to nearly 90%, suggesting a selective depletion of MHC class I-proficient neoplastic cells. Notably, the addition of mAb/IL2 to ICB almost completely prevented the outgrowth of MHC class I-deficient cells, augmented the tumor rejection rate to 60%, and significantly improved survival.

Moreover, we found that mAb/IL2 also sensitizes MHC class I-deficient tumors for chemotherapy. In the MC38-Her2-*B2m^-/-^*model all tumor-bearing mice treated with either oxaliplatin (OX) or mAb/IL2 alone experienced tumor progression, while more than half of the mice treated with the combination of chemotherapy and mAb/IL2 experienced full tumor rejection and long-term survival (Fig. 2F and fig. S6I). Similarly, in the B16F10-*B2m^-/-^* model, tumors were unresponsive to cyclophosphamide (CTX) treatment, while adding mAb/IL2 to the chemotherapy rescued half of the mice by complete tumor rejection (Fig. 2G and fig. S6J).

In summary, our data show that MHC I deficient tumors respond to IL2 treatment when combined with mAb and that mAb/IL2 overcomes the general therapeutic resistance of MHC class I-deficient tumors and enables complete tumor rejections.

### IL2 treatment is associated with immune cell infiltration and an IFN-associated inflammatory signature in the TME of MHC class I-deficient tumors

To study the mechanisms that reverse the treatment resistance, we analyzed B16F10-*B2m^-/-^* tumors collected from the different treatment groups (Fig. 3A and fig. S7, A and B). In both control and mAb monotherapy-treated mice, tumors exhibited scarce CD45^+^ leukocyte infiltration, while IL2 treatment effectively reversed the TME desertification and reestablished immune cell infiltration (Fig. 3B). Tumors of mice treated with either IL2 alone or in combination with mAb were briskly infiltrated by various NK- and T-cell subsets and macrophages, while the infiltration by cDC1 was not significantly elevated (Fig. S8A). The infiltration by regulatory T cells (Treg) increased upon treatment, whereas the intratumoral CD8/Treg ratio remained high (Fig. S8B), owing to the pro-found concurrent CD8^+^ T-cell expansion.

**Figure 3.**
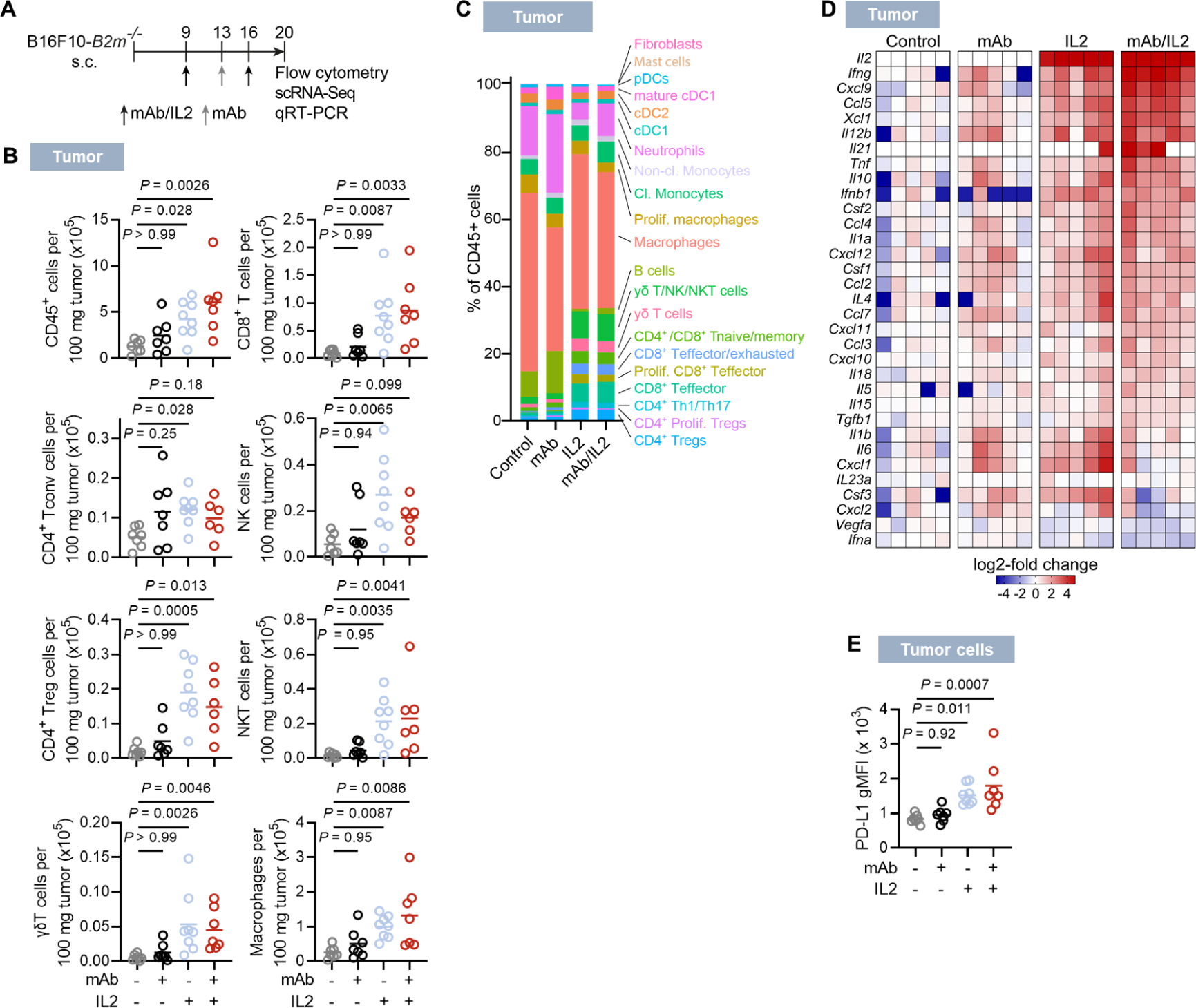
IL2 provokes an IFN-associated inflammatory signature in the TME. (**A**) B16F10- *B2m^-/-^* tumor model: Treatment and analysis scheme. (**B** to **E**) Mice were treated with αTrp1 mAb and intravenous IL2 mRNA-lipopolyplex (short IL2), either of the single compounds or isotype and albumin-encoding mRNA as controls initiated at day 9 after inoculation and tumors were analyzed 20 d after inoculation for infiltration of indicated cell subsets (B), proportions of cell type clusters across treatment groups as identified by single-cell RNA sequencing (scRNA-Seq) (C) expression of cytokines and chemokines as determined by qRT-PCR on mRNA isolated from whole-tumor tissues, depicted as log2-fold change over the control group (D) and PD-L1 expression in the tumor cells (E). Single outliers were removed by Grubb’s test (B). n=7 (mAb/IL2), n=7 (mAb), n=8 (IL2), n=7 (control) (B and E), n=3 pooled per group (C) and n=5 with each column representing an individual tumor (D).

Infiltrating immune cells displayed increased proportions of activated CD25^+^ CD4^+^ Tconv cells, CD25^+^ CD8^+^ T cells and KLRG1^+^ NK cells (Fig. S8C). Comparing the overall cellular composition of TILs, we found clusters of T-cell and NK-cell subsets expanded in IL2- and mAb/IL2- treated mice, while the relative proportion of neutrophils and B cells decreased (Fig. 3C and fig. S7A).

Gene expression analysis verified that IL2 mRNA treatment alone is sufficient to establish a strong inflammation signature with pronounced upregulation of *Ifng* and *Il2,* as well as of chemokines that recruit T cells and NK cells (*Cxcl9, Cxcl10*) and macrophages (*Csf1*, *Ccl2*) (Fig. 3D). PD-L1 expression was upregulated on tumor cells, indicating inflammatory signaling (Fig. 3E). The mAb/IL2 combination resulted in a similar molecular signature, yet with a substantially more enhanced expression of key genes such as *Ifng, Cxcl9, Il12b*.

In aggregate, these data show that IL2 is the main driver for igniting an inflammatory TME immune contexture that can be further enhanced by mAb, while mAb treatment alone has no significant impact on the inflammatory environment in *B2m^-/-^* tumors, which explains its lack of efficacy as a monotherapy.

### Activated CD8^+^ T cells and macrophages have a central role in rejection of MHC class I- deficient tumors in mAb/IL2-treated mice

In order to understand the mechanism underlying reversion of therapy resistance of MHC class I loss tumors by mAb/IL2, we conducted a series of *in vivo* cell depletion and cytokine neutralization experiments in mAb/IL2-treated mice bearing B16F10-*B2m^-/-^* tumors (Fig. S9, A and B). The efficacy of the mAb/IL2 treatment remained unaffected when either CD4^+^ T cells or neutrophils were depleted (Fig. 4, A and B and fig. S9, C and D).

**Figure 4.**
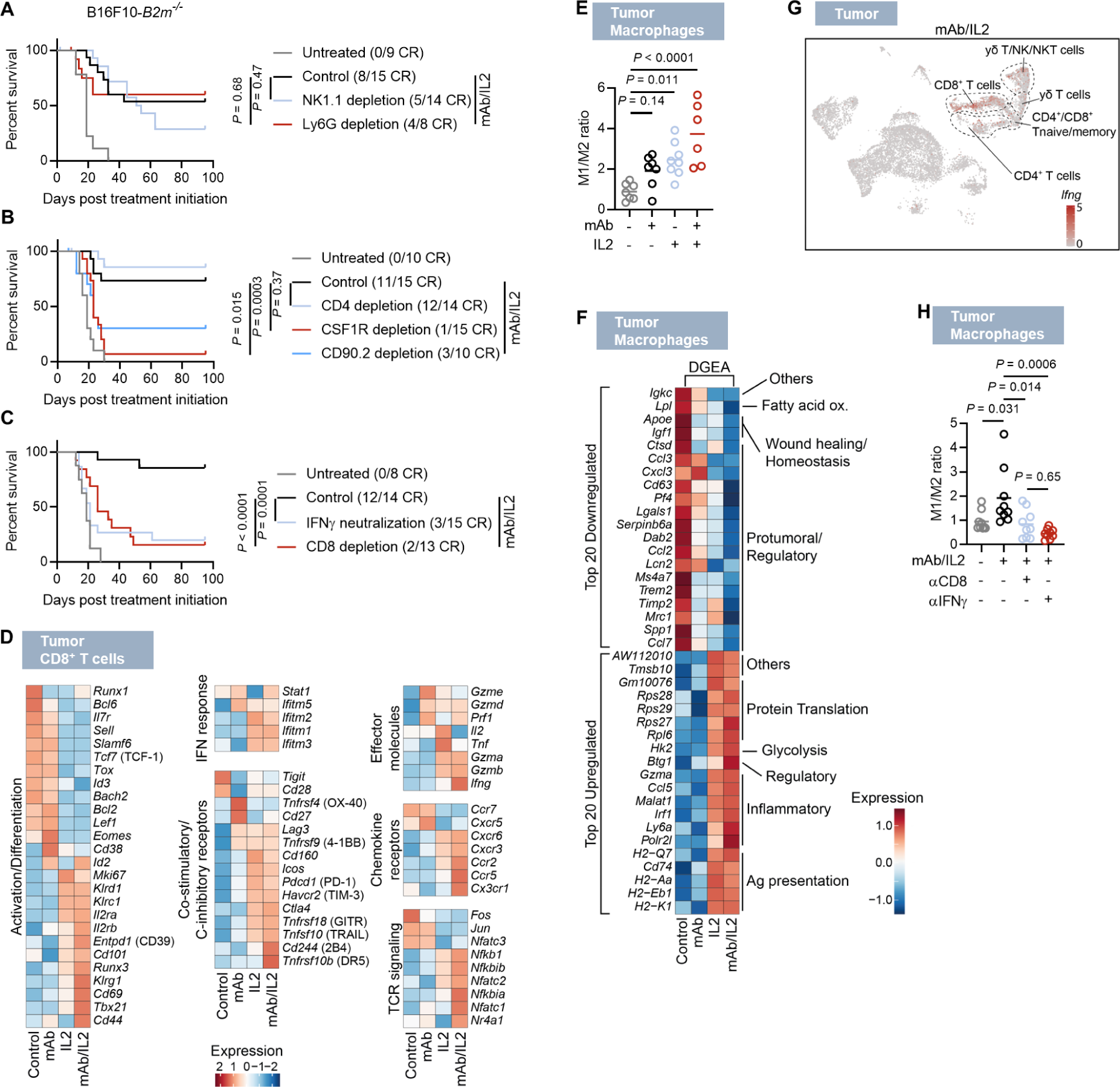
CD8^+^ T cells and macrophages have a central role in rejection of MHC class I- deficient tumors in mAb/IL2-treated mice. (**A** to **C**) Survival of mice treated with αTrp1 mAb and intravenous IL2 mRNA-lipopolyplex (short IL2) and depleted of specific immune cell subsets. Depleting/neutralizing mAbs were injected against NK1.1 and Ly6G (A), CD4, CSF1R and CD90.2 (B) or IFNγ and CD8 (C). Control groups were injected with an irrelevant mAb. (**D** to **H**) B16F10-*B2m^-/-^* tumor-bearing mice were treated with αTrp1 mAb and IL2 mRNA, either of the single compounds or isotype control and albumin-encoding control mRNA at day 9 after inoculation. 18-20 d after inoculation, tumors were analyzed for expression of selected genes in combined clusters containing CD8^+^ T cells determined by scRNA-Seq (D), M1/M2 ratio of infiltrating macrophages (E), the top 20 up-or downregulated genes in macrophages infiltrating mAb/IL2-treated tumors, compared to control by differential gene expression analysis (DGEA) using scRNA-Seq (F), expression of *Ifng* shown in a feature plot representing all leukocytes from mAb/IL2-treated tumors (G) and M1/M2 ratio of infiltrating macrophages after injection of depleting/neutralizing antibodies against CD8 or IFNγ or an irrelevant control mAb in mAb/IL2-treated mice (H). Single outliers were removed by Grubb’s test (E and H). n=8-15 (A). n=10-15 (B). n=8-15 (C). n=3 pooled per group (D, G and F). n=7-8 (E). n=9-10 (H).

Depletion of NK cells resulted in a minor loss in efficacy as it was associated with a statistically insignificant, slightly lower number of mice that experienced a complete response (Fig. 4A and fig. S9C). Depletion of CD8^+^ T cells (Fig. 4C and fig. S9E) or simultaneous depletion of all NK- and T-cell subsets (via CD90.2) (Fig. 4B and fig. S9D) resulted in lack of tumor rejection and a substantial decline in survival. Likewise, antibody-mediated neutralization of IFNγ (Fig. 4C and fig. S9E) and depletion of macrophages with CSFR1 antibodies (Fig. 4B and fig. S9D) almost completely abrogated the mAb/IL2-mediated treatment effect.

Having established CD8^+^ T cells and macrophages as essential mediators of the mAb/IL2 effect, we studied the molecular profiles of these cells. The transcriptome profiles of tumor-infiltrating CD8^+^ T cells in IL2- and mAb/IL2-treated mice showed upregulation of *Tbx21* and *Klrg1* as well as various effector molecules (e. g. *Ifng*, *Prf1*, *Gzmb*), co-inhibitory (e. g. *Pdcd1*, *Havcr2*) and costimulatory (e. g. *Tnfrsf9*, *Tnfrsf18*) receptors and IFN response genes (e. g. *Ifitm1*, *Ifitm2*) in line with a strong activation and differentiation of T cells into effector cells (Fig. 4D). IL2 administration alone or in combination with mAb induced expression of both *Il2ra* and *Il2rb*, indicating upregulation of the trimeric IL2 receptor that binds wild-type IL2 with high affinity. Again, changes in gene expression patterns were largely similar between IL2 or mAb/IL2-treated groups. Distinctive in mAb/IL2 treated mice was enhanced upregulation of genes involved in TCR signaling, e.g., *Ifng and Nr4a1* (32), indicating that cognate antigen encounter is augmented by addition of mAb to IL2 treatment.

Macrophages were polarized towards a proinflammatory M1-like phenotype in tumors treated with either IL2 or mAb/IL2 (Fig. 4E and fig. S10A). In mice treated with mAb/IL2, macrophages displayed a decrease in CD206, tumor-supporting immune regulators (such as *Ccl2, Pf4*), and an increase in the IFNγ-inducible markers PD-L1 and MHC class II (Fig. 4F and fig. S10, A and B). Moreover, macrophages showed high expression of T-cell attracting chemokines (*Ccl5*, *Cxcl9*, *Cxcl10*), inflammation-associated markers (e. g. *Irf1, Gzma*), tumoricidal mediators (*Nos2*) and genes related to antigen presentation (e. g. *H2-Q7, H2-Aa*), all in line with an M1-skewed phenotype (Fig. 4F and fig. S10, C and D). The co-stimulatory ligands CD80 and CD86 were highest in macrophages from the mAb/IL2 treated group, indicating the combined treatment led to superior T-cell stimulatory capacity.

Together, these data support that IL2 treatment is the main driver of the pro-inflammatory reprogramming of CD8^+^ T cells and macrophages, whereas on the transcriptome level additive effects of mAb are modest and associated with markers of augmented costimulatory capacity of macrophages and cognate antigen encounter of T cells.

### IL2-activated CD8^+^ T cells and macrophages cross-presenting tumor neoantigens engage in a cognate crosstalk

IFNγ was also flagged as critical for mAb/IL2 efficacy by our depletion studies (Fig. 4C and fig. S9E). Single-cell RNA-Sequencing (scRNA-Seq) analysis of all cell populations revealed effector CD8^+^ T cells as major source of the IFNγ production induced by mAb/IL2 (Fig 4G and fig. S10E). Both depletion of CD8^+^ T cells and neutralization of IFNγ prevented that macrophages resident in mAb/IL2-treated tumors acquired a M1-like phenotype (Fig. 4H), verifying the role of CD8^+^ T cell-released IFNγ in macrophage reprogramming.

As previously reported, M1 polarized macrophages have the capacity to eliminate antibody-opsonized tumor cells via antibody-dependent cellular phagocytosis (ADCP) after Fcγ receptor (FcγRs)-mediated uptake (33). We verified *in vitro* that IFNγ promotes the acquisition of an M1- like phenotype in bone marrow-derived macrophages (BMDM) (Fig. S10F), and IFNγ simulated BMDMs efficiently took up mAb-opsonized tumor cells (Fig. S10G). We found that in our model systems macrophages infiltrating *B2m^-/-^* tumors display the highest expression levels of Fcγ receptors (FcγRs) amongst all TILs (Fig. S10H), while γδT cells and NK cells, also potential antibody-sensing effector cells, exhibited only minute expression of FcγRs (Fig. S10I). Genes encoding FcγRs were further upregulated by treatment with IL2 with or without mAb in macrophages (Fig. S10J). In contrast, IL2 treatment failed to restore FcγR expression to relevant levels in γδT cells and NK cells (Fig. S10K).

These data indicate that IFNγ secreted by IL2-activated CD8^+^ T cells reprograms macrophages towards a phenotype capable of ADCP.

Macrophages are reported to be capable of cross-presenting antigens that they have engulfed in the TME (34). We found strongly enhanced expression of genes related to antigen presentation via MHC class I in *B2m^-/-^* tumors of mice treated with IL2 without or with mAb (Fig. 4F and fig. S10, C and D). As shown in mice with B16F10-*B2m^-/-^* tumors expressing Ovalbumin (Ova) containing the MHC class I-restricted SIINFEKL epitope (B16F10-Ova-*B2m^-/-^* cells), IL2 without and with mAb significantly enhanced the presentation of this antigen (Fig. 5A).

**Figure 5.**
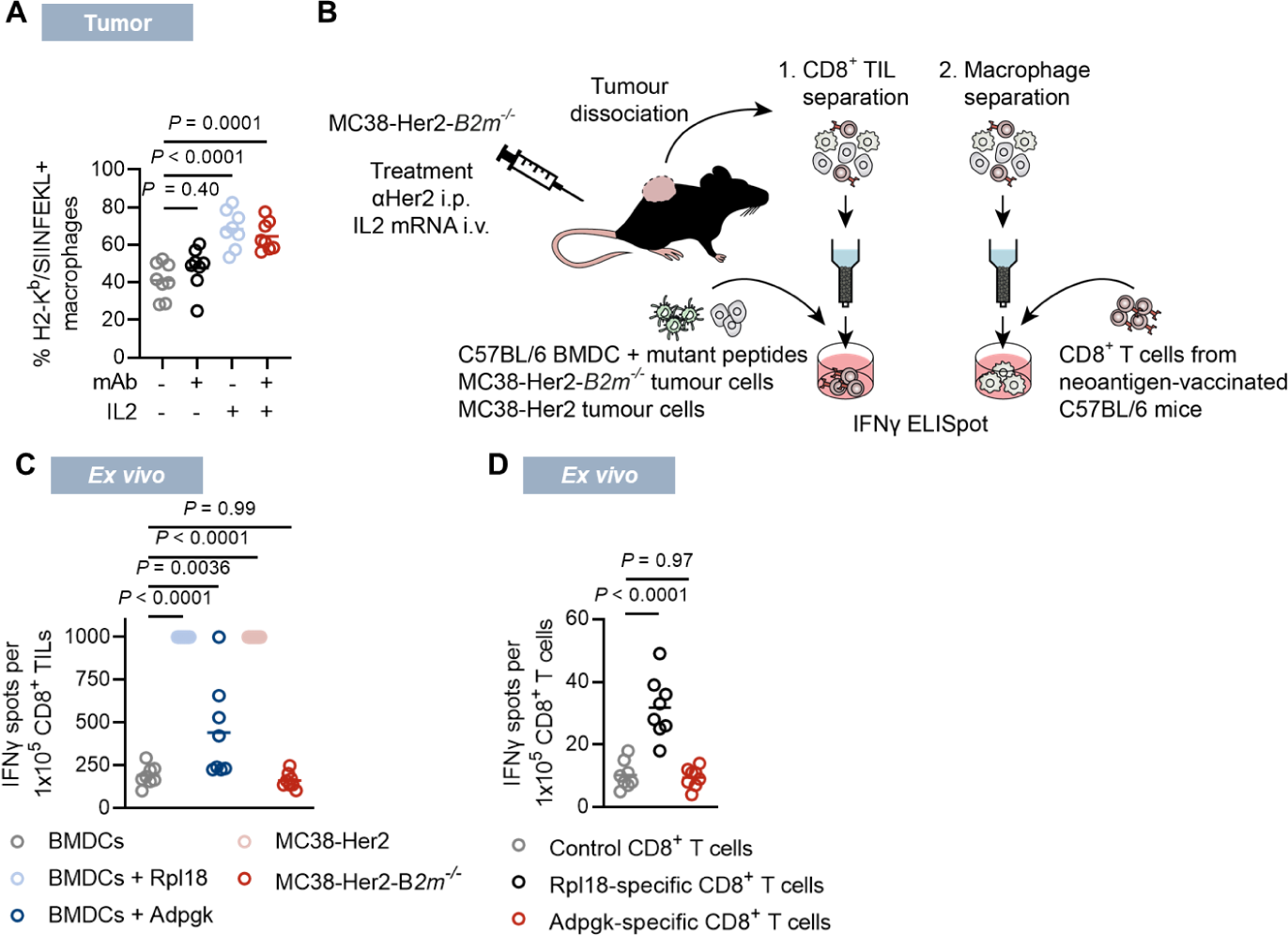
IL2-activated CD8^+^ T cells and macrophages cross-presenting tumor neoantigens engage in a cognate crosstalk. (**A**) B16F10-Ova-*B2m^-/-^* tumor-bearing mice were treated with αTrp1 mAb and IL2 mRNA, either of the single compounds or isotype control and albumin-encoding control mRNA at day 15 after inoculation. 18 d after inoculation, tumor-infiltrating macrophages were stained for H2-K^b^/SIINFEKL complexes. (**C** to **D**) MC38-Her2-B2m^-/-^ tumor-bearing mice were treated with αHer2 mAb/IL2 at day 9 after inoculation. 20 d after inoculation, tumors were harvested. Experimental scheme (B). CD8^+^ TILs were isolated and subjected to IFNγ ELISpot using BMDC or the indicated tumor cell lines (C) and tumor-infiltrating macrophages were isolated and subjected to IFNγ ELISpot with CD8^+^ T cells from mice immunized with the indicated RNA-LPX (D). n=8 (A, C, D).

To test whether uptake and presentation of tumor antigens by macrophages to CD8^+^ T cells in the TME may contribute to the rejection of *B2m^-/-^* tumors in mAb/IL2-treated mice, we made use of the MC38 model and of two of its mutations that act as MHC class I-restricted neoepitopes driving the strong immunogenicity of this tumor. The immunodominant ribosomal protein L18 (Rpl18)- derived neoepitope is highly expressed in MC38 (Fig. S10L) and Rpl18 neoantigen-specific CD8^+^ T cells are known to contribute significantly to the immunological control of MC38 tumor growth in ICB-treated mice (35). The other neoepitope is derived from a mutation in the ADP dependent Glucokinase (Adpgk) gene and exhibits low expression in MC38 (Fig. S10L). We isolated tumor-infiltrating macrophages and CD8^+^ T cells from MC38-Her2-*B2m^-/-^* tumor-bearing mice treated with mAb/IL2 and characterized them separately in a split-IFNγ ELISpot assay (Fig. 5B). The specificity of CD8^+^ T cells infiltrating mAb/IL2-treated tumors was assessed by co-culturing with bone marrow-derived DCs (BMDC) loaded with Rpl18 or Adpgk mutant peptides. While CD8^+^ T cells did not recognize MC38-Her2-*B2m^-/-^*cells, they released large amounts of IFNγ when cultured with Rpl18-pulsed BMDC and lower amounts with Adpgk-pulsed target cells (Fig. 5C), indicating that even though tumors were MHC class I-deficient, CD8^+^ T cells directed against tumor neoantigens were prevalent in the TME. In a complementary experimental setting, macrophages isolated from mAb/IL2-treated mice were co-cultured with Rpl18- or Adpgk-specific CD8^+^ T cells obtained from mice immunized with mRNA encoding these mutant neoantigens (Fig. S10M). Strong IFNγ release by Rpl18-specific CD8^+^ T cells (Fig. 5D) suggested that macrophages in the TME had captured and processed tumor-expressed Rpl18 *in situ* for cross-presentation.

## Discussion

In summary, our study provides a number of important findings that have implications for the understanding and treatment of tumors with defective MHC class I antigen presentation.

First, we show that MHC class I loss not only renders cancer cells resistant to CD8^+^ T-cell recognition and killing, but also leads to a secondary immune desertification of the TME, with loss of the IFNγ-response signature inherent to inflamed tumors, decrease of chemokines and of IL2 expression and exclusion of inflammatory immune cells, a phenotype similar to that of poorly CD8^+^ T-cell infiltrated and MHC class I-deficient areas of human tumors (6,7,36,37).

Second, we show that MHC class I-deficient cancers exhibit a general therapeutic resistance encompassing lack of response to immune-, chemo-and radiotherapy. Our finding is consistent with previous reports describing the contribution of T cell adaptive immunity to the therapeutic effect of these treatment modalities (38–41), and provides a mechanistic understanding for clinical studies linking decreased MHC class I expression to resistance to chemotherapy (42). The low frequency of tumor antigen-specific CD8^+^ T cells we detected after radiotherapy of MHC class I- deficient tumors suggests not only compromised T-cell effector function but also impaired T-cell priming upon tumor antigen release in tumors lacking functional MHC class I expression. This most likely results from decreased infiltration by cDC1, which intercept and cross-present tumor antigens released upon irradiation or other cytotoxic treatments and thus trigger CD8^+^ T-cell responses (43).

Third, we found that administration of an mRNA-encoded Interleukin-2 (IL2) reinstates an IFNγ- response signature and an inflamed and immune cell infiltrated TME contexture. When combined with a tumor-targeting monoclonal antibody (mAb), this successfully overcomes therapeutic resistance of MHC class I-deficient tumors and leads to complete tumor rejections in mouse tumor models.

We delivered IL2 as an mRNA-lipopolyplex, encoding an Albumin-IL2 fusion protein, that is administered intravenously and then taken up and translated in the liver. We have shown for this platform that the Albumin-IL2 fusion protein is secreted into the circulation, has an extended half-life in serum, and accumulates in the tumor (Peters *et al.*, manuscript submitted), thus resulting in prolonged cytokine exposure of immune cells in the TME at concentrations that are not associated with toxic effects. The prolonged activity of our IL2 format helps to overcome the pharmacological limitations of recombinant IL2 that may have compromised the therapeutic potential of this inflammatory cytokine in previous clinical trials testing the mAb/IL2 combination therapy (44,45). The IND-supporting safety data of this Albumin-IL2 fusion encoding mRNA showed tolerability at biologically active dose levels in non-human primate PK/PD studies and safety in mouse GLP toxicity studies.

Finally, and most importantly, our study reveals an unexpected mechanism by which tumor-antigen-specific CD8^+^ T cells drive antitumor activity against MHC class I-deficient tumors upon mAb/IL2 treatment.

The role of CD8^+^ T cells is counterintuitive, also given that previous studies reported NK cells as mediators of MHC class I-deficient tumor control. While those studies were undertaken with recombinant IL2 variants engineered for reduced binding to the high-affinity trimeric IL2 receptor (24,25), a recent study has demonstrated that binding to the high-affinity complex including IL2 receptor α (CD25) is critical for the expansion of antigen-specific CD8^+^ T cells (26). We chose to use wild-type IL2 with preferential binding to CD25, which is upregulated on CD8^+^ T cells upon cognate antigen encounter. This may explain why our IL2 format combined with tumor-targeting mAbs induces superior tumor rejection with only minor contribution of NK cells.

Our data show that IL2 treatment together with IFNγ secreted largely by activated CD8^+^ T cells promote an inflamed and infiltrated TME. Tumor-resident macrophages upon IL2 treatment undergo M1-type reprogramming with profound transcriptomic changes. The M1 phenotype is known to be associated with augmented capabilities to take up and eliminate tumor cells via ADCP (33), to directly kill cells via release of tumoricidal compounds (46), recruit T cells (47), cross-present antigen to and activate T cells (48). In our experiments, adding the mAb to IL2 increases TCR signaling and IFNγ production by CD8^+^ T cells, suggesting that the mAb may increase the capacity of macrophages to take up and cross-present antigens.

Our data with cells isolated from *B2m*^-/-^ tumors confirms functionally that resident macrophages are indeed capable to present cancer antigens and infiltrating CD8^+^ T cells are able to recognize those. In response to encountering macrophage-presented cancer antigens, CD8^+^ T cells become the major source for high IFNγ secretion into the TME, via which they are according to our depletion experiments together with macrophages pivotal for mAb/IL2-mediated tumor rejection. IFNγ is a pleiotropic cytokine known for its potent promotion of an inflammatory tumor micromilieu. IFNγ has direct tumoricidal activity (49), suppresses angiogenesis (50), and inhibits cancer cell proliferation (51). Previous reports show that bystander eradication of antigen-loss variant tumor cells depend on tumor antigen presentation by stromal cells and IFNγ released by CD8^+^ T cells (31), exemplifying the anti-tumor effect exerted by IFNγ.

Most likely various mechanisms contribute ultimately to tumor cell killing and tumor rejection in our mouse models. The observation that for clinical efficacy of prolonging mouse survival, tumor rejection and reversion of treatment resistance, IL2, that drives the major TME changes, is not sufficient but needs to be combined with mAb may indicate that for ultimate tumor eradication Fc-mediated mechanisms such as ADCP may have a prominent role.

Besides macrophages, cDC1 are considered the most effective cross-presenting cell type and are specialized in activating CD8^+^ T-cells (52). Although the infiltration of cDC1 was very low in MHC class I-deficient tumors in the first place and mAb/IL2 did not significantly elevate their infiltration into the tumors, we cannot exclude a potential role of these cells in fostering the CD8^+^ T-cell response.

Whereas mAb/IL2 treated mice develop immunological memory against B16F10-*B2m^-/-^* tumors our re-challenge experiments demonstrate that memory is insufficient to fully protect from tumor re-growth. At the time of re-challenge, all administered mAb can be expected to be eliminated from the system and without an inflammatory stimulus, immunogenicity of B16F10-*B2m^-/-^* is probably too low to recall a protective T-cell response.

One potential implication of our research is that combining mAb/IL2 with anti-PD-1 or other immune checkpoint inhibitors, could further enhance their therapeutic efficacy. Antagonizing PD-1 expression on macrophages is reported to promote mechanisms that synergize with those we have observed under mAb/IL2 treatment, namely phagocytosis (53), M1-polarization (54) and differentiation of myeloid progenitors into inflammatory macrophages (55).

In conclusion, our study highlights the unexpected significance of restoring neoantigen-specific T cell immunity in cancers with MHC class I deficiencies that are strongly activated by macrophage cross-presented tumor antigens to promote an inflammatory tumor environment counteracting tumor-cell intrinsic resistance mechanisms.

In summary, our findings offer new mechanistic insights and strategies to address the increasing medical need for treating immunotherapy resistant cancers characterized by MHC class I loss and immune desertification.

## Acknowledgments

We thank I. Beulshausen, E. Petscherskich, E. Russell, S. Zenner, A. Selmi, L. Cao, M. Brkic, Y. Feuchter, C. Born, L. Kögler, J. Dallmann and S. Silakhori for excellent technical assistance throughout the *in vivo* studies; U. Schmitt for dedicated support in the cell culture; S. Witzel for planning and Z. Yildiz and U. Schmitt for cloning of lentiviral constructs; M. Perkovic and J. Kaipan for producing lentiviral supernatants and performing transductions; S. Gangi Maurici and N. Köhl for assisting the design and for performing qRT-PCR assays; Ö. AkilliÖztürk and C. Leber for histological analysis of tumor tissue; D. Schneider and K. Wolf for performing clinical chemistry; S. Schön and T. Franze for technical assistance in histopathology; D. Eisel for assisting the cell sorting; N. Correia and T. Bukur for supporting scRNA-Seq. We are grateful to L.M. Kranz and N. Salomon for fruitful discussion. We thank Andrew Finlayson for providing medical writing support and editorial support, in accordance with Good Publication Practice (GPP3) guidelines.

## Funding

This work was supported by the European Research Council Advanced Grant “SUMMIT” (789256 to U.S.).

## Author contributions

M. Vormehr and U. Sahin conceptualized the work. J.D. Beck, M. Vormehr, M. Diken, S. Kreiter and U. Sahin planned and analyzed experiments. J.D. Beck and M. Vormehr performed experiments. J.D. Beck, M. Vormehr, M. Diken, S. Kreiter, Ö. Türeci and U. Sahin interpreted the data. J.D. Beck, M. Vormehr, Ö. Türeci and U. Sahin wrote the manuscript. M. Suchan designed and analyzed qRT-PCR assays. M. Streuber, E. Diken and L. Kolb performed scRNA-Seq. L. Allnoch performed histopathological assessments. F. Vascotto performed in vitro ADCP assays. D. Peters performed the PK analysis. T. Beißert established CRISPR/Cas9 protocols.

## Declaration of interests

J.D. Beck, M. Vormehr, L. Allnoch, M. Diken and S. Kreiter are employees and Ö. Türeci and U. Sahin are cofounders and management board members at BioNTech SE (Mainz, Germany). J.D. Beck, M. Vormehr, L. Allnoch, M. Diken, S. Kreiter, Ö. Türeci and U. Sahin hold securities from BioNTech SE. J.D. Beck, M. Vormehr, M. Diken, S. Kreiter and U. Sahin are inventors on patents or patent applications related to this study.

## Data and materials availability

The data that support the findings of this study as well as custom code are available from the corresponding author upon reasonable request.

## Materials and Methods

### Mice

C57BL/6 mice were purchased from Envigo, BALB/c mice from Janvier. 9-12 weeks old age-matched female mice were used throughout the studies and maintained in accordance with federal and state policies on animal research in a specific-pathogen-free facility at BioNTech SE. Group sizes and experimental procedures were approved by regulatory state authorities for animal welfare.

### Tumor cell lines

CT26 colon carcinoma and B16F10 melanoma cell lines were purchased from ATCC (CT26: CRL-2638, lot no. 58494154; B16F10: CRL-6475, lot no. 58078634). MC38 colon adenocarcinoma cells were kindly provided by F. Ossendorp (Leiden University). Master and working cell banks were generated upon receipt and cells were passaged not more than eight times before use in in vivo experiments. Cells were cultured in DMEM (Life technologies) supplemented with 10% FBS (B16F10), RPMI1640 (Life technologies) supplemented with 10% FBS (Sigma) (CT26) or RPMI1640 (ATCC) supplemented with 10% FBS and 2 mM L-Gln (MC38) and 4 µg/mL Blasticidin (InvivoGen) (MC38-Her2-*B2m^-^*^/-^). Primary immune cells were kept in RPMI1640 medium (Life technologies) supplemented with 10% heat-inactivated FCS, 1 mM sodium pyruvate (Life technologies), 1x MEM Non-Essential Amino Acids Solution (Life technologies) and 50 U/mL Pen/Strep (Life technologies). All cell lines were regularly confirmed to be free of mycoplasma contaminations.

### Generation of stable transgene-expressing cell lines

Codon-optimized rat Her2/neu (UniProt accession number P06494) was synthesized by GeneArt (Thermo Fisher Scientific) and the manually codon-optimized chicken Ovalbumin gene (UniProt accession number P01012) was synthesized by Eurofins. In brief and as previously described (56), the target genes were inserted into the lentiviral backbone pLenti6.4/R4R2/V5-DEST multisite Gateway destination vector under the control of the human EF1a promoter. Lentiviral particles were produced in HEK293T cells by cotransfection of the lentiviral vector together with pCMVDR8.91 and M620. The viral particles were loaded onto non-treated cell culture plates (Corning) coated with Retronectin (Takahara) and target cells were plated and cultured for 3 days. Subsequently, cells were transferred to selective medium containing 4 µg/mL Blasticidin (Invivogen) (MC38-*B2m^-/-^* transduced with Her2/neu) or 300 µg/mL Hygromycin (Invitrogen) (B16F10-*B2m^-/-^* transduced with Ovalbumin) and cultured for 7 days. Single clones were obtained by limiting dilution. Single clones with stable integration of the transgene as shown by retained expression after culturing the cells in non-selective medium for 4 weeks was selected for further usage.

### Generation of knockout cell lines

The CRISPR/Cas9 system was used for genetic knockouts. sgRNA sequences targeting *B2m* (3’-CATGGCTCGCTCGGTGACCC-5’ for CT26 and B16F10 and 3’-GACAAGCACCAGAAAGACCA-5’ for MC38) were generated using publicly available tools (E-CRISP, CCTop or CRISPRko). sgRNAs and mRNA encoding for Cas9 were delivered to cells via electroporation or lipofection (RNAiMAX, Invitrogen) together with mRNAs encoding for the viral proteins E3 and B18 in order to inhibit a cellular response to dsRNA. Single clones were obtained by limiting dilution, expanded and screened by flow cytometry and IFNγ ELISpot for cell surface expression of MHC class I and recognition by tumor antigen-specific CD8+ T cells as proxies for *B2m* deletions. For each knockout cell line, several clones were tested in vitro and clones with confirmed MHC class I loss that resembled the parental cells with regards to morphology and in vitro growth behavior were selected for further experiments and cell banks were generated.

### Tumor models

For the inoculation of tumors, tumor cells (5×10^5^ CT26/CT26-*B2m^-^*^/-^ in BALB/c mice, 5×10^5^ MC38/MC38-*B2m^-^*^/-^/MC38-Her2-*B2m^-^*^/-^ and 3×10^5^ B16F10/B16F10-*B2m^-^*^/-^ in C57BL/6 mice) suspended in 100 µL sterile PBS were injected s.c. into the right flank of the mice. For early treatment models, in which treatments were initiated 5 days or less after tumor inoculation, mice were randomly assigned to treatment groups once tumors were palpable (not more than 1 mm³ in size). In the remaining experiments, mice were allocated into treatment groups based upon their tumor sizes using the ANOVA P method implemented in Daniel’s XL tool box (v7.3.4). Tumor diameters were measured 2-3x /week with a caliper and the tumor size was calculated using the formula (A x B2)/2 with A as the largest and B as the smallest diameter. Mice were sacrificed and recorded as dead when they reached defined endpoint criteria, including a tumor size of 1500 mm³ or over, tumor ulceration, or signs of distress or moribundity. Re-challenge was performed by s.c. injection of 3×10^5^ B16F10-*B2m^-^*^/-^ cells into the contralateral flank of mAb/IL2 treated mice cured from B16F10-*B2m^-^*^/-^ tumors at day 72 of the experiment.

### Immune checkpoint blockade

Treatment of B16F10/B16F10-*B2m^-^*^/-^ with αPD-1 and αCTLA4 was initiated 3 days after tumor inoculation. Mice were treated with αPD-1 (clone RMP1-14, BioXCell) and αCTLA4 (clone 9H10, BioXCell) with a total of five i.p. injections every 3-4 days at doses of 200 µg or 100 µg, respectively. For the first injection, 200 µg αCTLA4 was applied. The control group received isotype controls (clone 2A3 and Syrian hamster IgG, BioXCell). Treatment of MC38/MC38-*B2m^-^*^/-^ with ICB was initiated when tumors reached a mean size of ca. 32-36 mm³. Mice received up to 6 i.p. injections every 3-4 days with 200 µg irrelevant antibody (clone 2A3, BioXCell), αPD-1 (clone RMP1-14, BioXCell) or αCTLA4 antibody (clone 9D9, BioXCell).

### RNA-LPX vaccination

Design and generation of pharmacologically optimized antigen-encoding mRNA constructs have been described (57–59). gp70 mRNA encodes the H-2Ld restricted epitope AH1423-431 derived from the mouse leukaemia virus envelope glycoprotein 70 with a V/A amino acid substitution at position 5 (60). OVA mRNA encodes the H-2Kb restricted epitope OVA257- 264 derived from chicken ovalbumin. mRNAs against MC38 neoantigens encoded the H-2Db restricted neoantigen Rpl18 or the H-2Kb restricted neoantigen Adpgk. Antigen-encoding in vitro-transcribed mRNAs (BioNTech) were formulated with liposomes (BioNTech) to yield RNA-LPX as previously described24. Immunization of CT26 or CT26-*B2m^-^*^/-^ tumor-bearing mice was performed by injecting 40 µg of gp70 mRNA-LPX or empty-vector control mRNA-LPX r.o. on days 5, 8, 12 and 19 after tumor inoculation. In order to yield antigen-specific CD8^+^ T cells for ELISpot assays, 2-3 weekly immunizations with 20-40 µg mRNA were performed. BALB/c mice were immunized with gp70 RNA-LPX and C57BL/6 mice were immunized with OVA, Rpl18 or Adpgk RNA-LPX.

### Chemo-and radiotherapy

MC38/MC38-*B2m^-^*^/-^ tumor-bearing mice were injected i.p. with 5 mg/kg OX (Medac) in 5% Glucose (Braun) on days 4, 11 and 18 after tumor inoculation. CT26/CT26-*B2m^-^*^/-^ tumor-bearing mice were treated with a combination chemotherapy consisting of 5 mg/kg OX injected i.p. and 60 mg/kg 5-fluorouracil (5-FU) (Medac) in 0.9% NaCl (Braun) injected r.o.. Treatment of CT26/CT26-*B2m^-^*^/-^ tumors was initiated when the different cohorts reached mean tumor sizes of approximately 6 mm³. Three weekly cycles were performed in total. Cyclophosphamide (CTX) supplied by Baxter was injected i.p. into B16F10-*B2m^-^*^/-^ tumor-bearing mice at a single dose of 150 mg/kg in 0.9% NaCl 8 days after inoculation.

To facilitate the LRT of tumors while shielding the rest of the animal, a custom-made lead shield was used that contains 1.5 cm wide circular cavities under which the tumors were positioned. Mice were irradiated with a single dose of 12 Gy using an X-RAD 320 (Precision X-Ray Instruments) when the different cohorts of CT26/CT26-*B2m^-^*^/-^ tumor-bearing mice reached mean tumor sizes of approximately 30 mm³.

### mAb/IL2 therapy

mRNA encoding mouse IL2 (positions 21-169) (GenBank X01772.1) fused to the C-terminus of mouse albumin (GenBank AK050248.1) via a linker (G2SG4SG2) was generated from a DNA template by in vitro transcription in the presence of the trinucleotide cap 1 analogue ((m27,3’-O)Gppp(m2’-O)ApG; TriLink) and all four nucleotides with 1-methylpseudouridine-5’-triphosphate (m1ΨTP; Thermo Fisher Scientific) substituting for uridine-5’-triphosphate (UTP) (61). Non-coding backbone sequence elements were designed for improved mRNA stability and translational efficiency and included a 5’ UTR derived from Tobacco Etch Virus (TEV) genomic RNA leader and a 3’ UTR comprising two sequence elements derived from the amino terminal enhancer of split (AES) mRNA and the mitochondrial encoded 12S ribosomal RNA (62) and a poly(A) tail (100 nucleotides) interrupted by a linker (A30LA70, 10 nucleotides) (57). Double-stranded mRNA contaminants were depleted by cellulose purification (63). The mRNA was formulated with a polymer/lipid-based transfection reagent (TransIT-mRNA Transfection kit (Mirus Bio)) to obtain mRNA-lipopolyplexes that target to the liver upon intravenous administration (29). Batches of formulated mRNA were prepared for up to five injections at once. Per injection, 1 µg mRNA in 1 µL H2O was added to 97.16 µL ice-cold DMEM, followed by 1.12 µL TransIT-mRNA Reagent and 0.72 µL mRNA Boost Reagent. The mixture was incubated for 2 min at room temperature and injected intravenously once a week for up to four times. mAbs against Trp1 (clone TA99, BioXCell) or Her2 (clone C1.18.4, BioXCell) were injected i.p. at doses of 200 µg. mRNA encoding albumin and an IgG2a isotype control mAb (clone C1.18.4, BioXCell) served as controls. The mAbs were injected every 3-4 days for up to seven times.

### Tissue preparation and cell isolation

Single-cell suspensions from tumors were prepared using the Tumor Dissociation Kit, mouse (Miltenyi Biotec) in combination with a gentleMACS^TM^ Dis-sociator (Miltenyi Biotec) according to the manufacturer’s instructions. In brief, tumors were harvested, transferred to gentleMACSTM C Tubes (Miltenyi Biotec) prefilled with 2.5 mL of the enzyme mix prepared in RPMI1640. Tumors were roughly chopped into pieces and digested using the preprogramed protocols for subcutaneous mouse tumors. The resulting cell suspensions were mashed through 70 µM cell strainers (Greiner Bio-One) while rinsing with RPMI1640. Cells were collected by centrifugation (6 min, 460 x g) and erythrocytes were lysed by incubating the cell suspension for 5 min in 5 mL of a hypotonic electrolyte solution. Afterwards, 15 mL PBS containing 2 mM EDTA were added, and the cell suspensions were transferred through a 70 µM cell strainer in order to remove aggregates of lysed erythrocytes. The cells were washed with 10 mL PBS containing 2 mM EDTA, resuspended in medium after a final centrifugation step and stored at 4 °C until being used for flow cytometry staining. To generate single-cell suspensions from lymph nodes, lymph nodes were manually minced into pieces and digested with collagenase A (1 mg/mL; Roche) and DNAse (10 µg/mL; Sigma) in RPMI1640 for 10 min at 37 °C and the resulting suspensions were transferred through 70 µM cell strainers, which were rinsed using RPMI1640. Single-cell suspensions from spleens were prepared by mashing the tissues through 70 µM cell strainers followed by hypotonic lysis to remove erythrocytes. Cells were collected by centrifugation and stored as describe above. Tumor-infiltrating macrophages and CD8^+^ T cells were isolated by magnetic-activated cell sorting (MACS) using Anti-F4/80 MicroBeads UltraPure, mouse (Miltenyi Biotec) or CD8 (TIL) MicroBeads, mouse (Miltenyi Biotec), respectively, and CD8^+^ T cells were isolated from splenocytes using CD8a (Ly-2) MicroBeads, mouse (Miltenyi Biotec), according to the manufacturer’s instructions.

### IFNγ Enzyme-linked ImmunoSpot

**(ELISpot)** IFNγ ELISpot assays to detect the release of IFNγ by T cells upon recognition of antigen was performed as described23. To validate absence of MHC class I in tumor cells after knock-out of *B2m*, 5×10^5^ splenocytes from immunized mice were co-cultured with 5×10^4^ tumor cells. Splenocytes cultured without tumor cells served as background control. To detect antigen presentation by macrophages, 1×10^5^ CD8+ T cells isolated from spleens of RNA-LPX immunized mice were co-cultured with 5×10^4^ macrophages or with 5×10^4^ BMDCs and 2 µg/mL peptide as positive control. To analyze the antigen reactivity of CD8^+^ TILs, 1×10^5^ CD8+ TILs were co-cultured with 5×10^4^ tumor cells or 5×10^4^ BMDCs and 2 µg/mL peptide. BMDCs without peptide served as control. All co-cultures were performed for 18-24 h.

### Flow cytometry and fluorescence-activated cell sorting

**(FACS)** For analyses comparing the TME of untreated wild-type and *B2m^-^*^/-^ tumors, single-cell suspensions from tumors and TDLNs harvested 20 days after inoculation were used. For the analysis of αTrp1 mAb/IL2 treated B16F10- *B2m^-^*^/-^ tumors, the tumors were harvested 2 or 4 days after the second IL2 treatment (18 or 20 days after inoculation). For flow cytometry analysis, single-cell suspensions from processed tissues were directly subjected to staining. Prior to FACS of tumor-infiltrating leukocytes, CD45^+^ cells were pre-enriched using CD45 (TIL) MicroBeads, mouse (Miltenyi Biotec) according to the manufacturer’s protocol. For the staining procedure, cell suspensions were transferred to 96-well U-bottom plates (Corning). Dead cells were stained with Fixable Viability Dye (eBioscience) or Dead Cell Stain (Thermo Fisher Scientific) diluted in PBS for 15 min at 4 °C. For staining of extracellular antigens, PBS supplemented with 5% heat-inactivated FBS and 5 mM EDTA was used as washing and staining buffer. If no Fc receptors were stained, Fc blockade was performed prior to extracellular staining with Mouse BD Fc Block (BD). Cell surface antigens were stained for 30 min at 4 °C. Her2 was labelled with 20 µg/mL αHer2 mAb followed by staining with goat anti-mouse secondary antibody. For staining of intracellular antigens, cells were fixed and permeabilized using the Cytofix/Cytoperm kit (BD Pharmington) for cytoplasmic antigens or the FoxP3/Transcription Factor Staining Buffer Set (eBioscience) for transcriptions factors according to the manufacturer’s instructions. For peripheral blood samples, whole blood was stained and subsequently lysed using FACS lysing solution (BD Biosciences). In order to determine absolute cell counts, samples were transferred to Absolute Counting Tubes (BD). The number of cells/µL was calculated using the following formula: (# cell gate / # bead gate) x (beads per tube / sample V [µL]). Data was acquired using an LSRFortessa Flow Cytometer (BD) or FACSCanto II (BD) and FACSDiva Software (BD). A FACSMelody (BD) was used for cell sorting. Data analysis was performed using Flowjo (v10.4; Tree Star) software. See Supplementary Information for a full list of reagents used for staining.

### Immunohistochemistry

**(IHC)** For the analysis of immune cell infiltration, subcutaneous tumors were harvested from the mice, fixed with 4 % formalin (Carl Roth) overnight at 4 °C and embedded in paraffin blocks (FFPE). 3 μm tissue sections were stained with polyclonal goat anti-mouse CD8α (1:2000, Invitrogen) antibody. Briefly, after deparaffinization in xylene (Carl Roth) and rehydration in graded concentrations of ethanol in water, antigen retrieval using sodium citrate buffer (10 mM, pH 6) was performed for 15 min at 110 °C in a DAKO Pascal pressure chamber (Agilent). After blocking in PBS supplemented with 10% goat serum (Sigma Aldrich) for 30 min at room temperature sections were incubated with primary antibody overnight at 4 °C. Sections were washed in PBS, incubated in HRP-conjugated αRat secondary antibody (Abcam) for 30 min at room temperature and detection was performed with a streptavidin-HRP system (Vector Labs) in conjunction with counterstaining with Hemalum (Carl Roth). For immunofluorescence staining, after secondary antibody incubation, Opal 570 was used for signal amplification (1:100, Akoya Bioscience). Counterstaining was performed with DAPI for 5 min at room temperature (1 drop per 500 µL, Akoya Bioscience). Microscope images were taken using the Zeiss AxioImager M2 and analyzed using ZEN blue software (v2.3).

For the histopathological analysis of IL2 treatment effects, livers and lungs were harvested 5 days after treatment, fixed with 4 % formalin overnight at 4 °C and embedded in paraffin blocks (FFPE). 3 µm sections were stained with Modified Mayer’s Hematoxylin (Thermo Fisher Scientific) and Eosin Y (Thermo Fisher Scientific) using the ST5020 multistainer (Leica). Sections were scanned using a NanoZoomer S360 (Hamamatsu). Blinded histopathological assessment was performed by a veterinary pathologist.

### RNA In situ hybridization

**(ISH)** FFPE tissues were cut into 3-μm-thick sections. ISH was performed using a mouse NCR1 probe (Advanced Cell Diagnostics) and mRNAscope detection reagents 2.5 HD Brown (Advanced Cell Diagnostics) according to the manufacturer’s instructions. Standard pre-treatment was done using 1x Target retrieval buffer for 15 min at 95 °C (Advanced Cell Diagnostics) and Protease plus for 30 min at 40 °C. (Advanced Cell Diagnostics). Counter-staining was performed with Hemalum (Carl Roth) for 30 sec.

### qRT-PCR analysis

For comparing the cytokine and chemokine expression in wild-type and *B2m^-^*^/-^ tumors, the tumors were harvested 19 days after inoculation. For the analysis of αTrp1 mAb/IL2 treated B16F10-*B2m^-^*^/-^ tumors, the tumors were harvested 4 days after the second IL2 treatment (20 days after inoculation). Total mRNA was extracted from mouse tumors using the RNeasy Mini Kit (QIAGEN) and subsequently tested for concentration and integrity via NanoDrop 2000c (Thermo Fisher Scientific) and capillary gel electrophoresis using a Fragment Analyzer (Agilent Advanced Analytical). Reverse transcription was performed on 1 µg total mRNA per sample using PrimeScript™ RT Reagent Kit with gDNA Eraser (Takara) according to the manufacturer’s instruction. For the qRT-PCR analysis, SsoAdvanced™ Universal SYBR® Green Supermix (Bio-Rad) was used according to the manufacturer’s instructions. The qRT-PCR was performed in technical triplicates on a BioRad CFX384 Touch™ Real-Time PCR Detection System using 40 cycles of two-step PCR at an annealing and elongation temperature of either 60 °C or 62 °C depending on the primers and an annealing and elongation time of 30 sec. The reaction volume was 15 µl, including 1.25 µl cDNA per reaction and a final primer concentration of 333 nM per primer. To determine Ct-values the BioRad CFX Manager 3.1 software was used, calculation of relative expression values was performed using a ΔΔCt calculation. Normalization to Hprt was performed for assessment of Ifng and chemokine expression in normal versus *B2m^-^*^/-^ tumors and for comparison of the gene expression patterns in treated B16F10-*B2m^-^*^/-^ tumors, normalization to Hrpt and Tbp reference genes was performed. For reliability reasons a cut-off value, excluding Ct-values >35, was used. log2-fold changes of ΔΔCt values were plotted as heatmap using GraphPad PRISM 8. See Supplementary Information for a full list of primer sequences.

### Preparation of and sequencing of single-cell 3’-libraries for single-cell RNA-sequencing

**(scRNA-Seq)** B16F10-*B2m^-^*^/-^ tumors were used for scRNA-Seq 4 days after the second IL2 treatment (20 days after inoculation). Three biological replicates per treatment condition were barcoded with individual TotalSeq antibodies (Biolegend) and pooled in equal proportions. 2×10^4^ tumor-infiltrating leukocytes per sample were obtained by sorting viable CD45^+^ cells and were processed according to 10X Genomics Chromium Single Cell 3’ Reagent Guidelines (v3 chemistry with Feature Barcoding technology for Cell Surface Protein: CG000185 Rev C). In brief, ∼20,000 cells were loaded into each channel and partitioned into nanoliter-scale gel beads-in-emulsions using a 10X Chromium Controller. Cells were lysed and reverse transcription was performed using a C1000 TouchTM Thermal Cycler, followed by amplification and size selection to separate cDNA for 3’Gene Expression (GE) and Feature Barcoding (FB) library preparation. Enzymatic fragmentation was performed for GE libraries, whereas purification using Dynabeads™ MyOne™ Silane, attachment of adaptors and index sequences was done for both library types. Quantification of cDNA and final libraries were performed using the Qubit™ dsDNA HS Assay Kit and High-sensitivity DNA Chips. Final GE and FB libraries were diluted to 1 nM, pooled in a 16:1 ratio and loaded with a final concentration of 200 pM on two lanes of an Illumina NovaSeqTM 6000 S2 flowcell using the NovaSeqTM 6000 SP/S1/S2 SBS Cartridge (100 cycles) and NovaSeqTM 6000 SP/S1/S2 Buffer Cartridge. Libraries were sequenced using a NovaSeqTM 6000 with the following read lengths: 28 cycles (Read 1), 8 cycles (i7 Index) and 91 cycles (Read 2). This resulted in ∼38,000-58,000 reads per cell for GE libraries and ∼3,000-7,000 reads per cell for FB libraries.

### Processing of scRNA-Seq raw data

The 10X Genomics Cellranger single cell pipeline (v3.0.2) was used to perform sample demultiplexing, barcode processing and single-cell 3’ gene counting as previously described (64). Generated FASTQs were aligned to the mouse mm10 reference genome (v3.0.0) using the Cellranger count pipeline. Aligned reads were filtered for valid Unique Molecular Identifiers (UMIs) (max. 1 mismatch allowed) with sequencing quality score > 10 (90% base accuracy). Barcodes with UMI counts that fell within the 99th percentile of the range and PCR duplicates were removed. Principal component analysis (PCA) was followed to consider genes with at least one UMI count in at least one cell. The top 1,000 most variable genes were identified based on mean and dispersion. Technical replicates were combined to a single gene-barcode matrix containing all the data for each group using Cellranger aggr.

### Integration and clustering analysis of aggregated replicates

For integration and clustering analysis, the Seurat toolkit for R was used as described in the tutorials (https://satijalab.org/seurat/) (65). Low quality cells with less than 200 genes, more than three median absolute deviation (MADs) above the median mitochondrial genes and more than 3 MADs above the median gene counts were excluded from further analysis. Digital gene expression measurements and antibody-derived tags (ADTs) (FB data) were separately log-normalized and afterwards log-transformed. ADT files were used to identify singlets (cell barcodes with positive counts for one ADT), doublets (cell barcodes with positive counts for more than one ADT) or negatives (no ADT detected) (66). Highly variable genes (by default: 2,000) were identified for each dataset. Integration strategy was used to identify shared cell states (anchors) between multiple single-cell datasets and to harmonize them as previously described (67). A linear transformation step was applied to shift and scale the expression of each gene prior to a linear dimension reduction step via PCA using previously determined variable genes. Uniform-Manifold-Approximation and Projection (UMAP) (68,69) was performed prior to shared nearest neighbor graph-based clustering using the top 20 principal components (PCs) to identify cell clusters (70). A two-dimensional non-linear embedding of the cells was generated on the first 20 PC dimensions using UMAP. Wilcoxon rank-sum test (default) was applied to identify positive and negative markers of a single cluster compared to others clusters or all remaining cells via differential expression method. A set of canonical markers and differentially top expressed genes was used to identify populations of interest and assign each cluster with a cell type identity. A low-quality cell cluster (low number of genes/UMIs) and two doublet clusters were detected (co-expressing Cd3 and Cd14) and excluded from further analysis. For macrophages, neutrophils and B-cells, clusters were detected and combined to single clusters. A fibroblast cell population (co-expressing Sparc, Col1a1 and Col3a1) and a mast cell population (co-expressing Cpa3 and Gata2) were selected based on marker expression by the Seurat lasso tool. The final data set contained 52,794 cells.

### Mutation expression analysis

Quantification of neoepitope expression in MC38 cells based on whole-exome and RNA-sequencing data was performed as previously described (71). To determine the Mutation Expression, the gene transcript expression was multiplied with the allele frequency.

### Immune cell depletion and in vivo cytokine neutralization

Depletion of CD4^+^ T cells with αCD4 (clone YTS 191, BioXCell), CD8^+^ T cells with αCD8α (clone YTS 169.4, BioXCell) or total lymphocytes with αCD90.2 (clone 30H12, BioXCell) was performed by injecting 400 µg mAb 2 days before therapy initiation. Three more injections were performed 2, 5 and 9 (αCD4 and αCD90.2) or 12 (αCD8α) days after treatment initiation. Macrophages were depleted by injection of 600 µg αCSF1R (clone AFS98, BioXCell) 3 days before therapy. Five additional injections of 350 µg αCSF1R were performed every 2-3 days. Depletion of neutrophils with αLy6G (clone 1A8, BioXCell) or NK cells with αNK1.1 (clone PK136, BioXCell) was started 1 day before therapy by injecting 400 µg mAb. Five more injections of 200 µg mAb were performed 2, 6, 9, 13 and 15 days after therapy began. IFNγ was neutralized with αIFNγ (clone XMG1.2, BioXCell) by injecting 400 µg mAb 2 days before therapy and three more injections of 250 µg antibody 0, 2 and 5 days after therapy was initiated. In two experiments, clone 2A3 (BioXCell) was used as irrelevant antibody control and injected according to the scheme of αCD4 or αLy6G. For IFNγ neutralization and CD8 depletion studies, clone HRPN (BioXCell) was used as control and injected according to the scheme of αIFNγ.

### In vitro stimulation of BMDMs

Bone marrow cells were flushed from femur and tibia bones of C57BL/6 mice and homogenized by transfer through a 70 µm cell strainer. Erythrocytes were lysed by hypotonic lysis. 2×10^7^ cells were seeded into T75 culture flasks (Greiner) and cultured in the presence of 100 ng/mL recombinant mouse M-CSF (Peprotech) for 8 days. Adherent cells were harvested and macrophage phenotype was confirmed by microscopy and flow cytometry (CD11c-^¬^ GR-1^-^ CD11b^+^ F4/80^+^). For cytokine stimulation, 2×10^5^ BMDMs were seeded into 24-well plates and cultured for 24 h in the presence of 50 ng/mL IFNγ (Peprotech) and 100-1000 U/mL IL2 (Proleukin S, Novartis). BMDMs were detached by incubating in PBS + 10 mM EDTA for 10 min at 4 °C and subjected to flow cytometry analysis.

### In vitro ADCP

MC38-Her2-*B2m^-^*^/-^ cells were electroporated with mRNA encoding ZsGreen. After culturing the cells for 24 h, tumor cell opsonization was performed in medium containing 5 µg/mL αHer2 mAb or isotype control mAb for 10 min at 37 °C. 105 opsonized tumor cells were co-cultured with IFNγ-stimulated BMDMs at an E:T ration of 1:1 for 2 h at 37 °C. Subsequently, BMDMs were subjected to flow cytometry analysis to detect incorporated ZsGreen as a measure for tumor cell phagocytosis.

### Data representation, exclusion and statistics

The sample size (n) represents the number of biological replicates. All data are shown as mean and error bars represent SEM. Samples were excluded from the analysis if the number of cells obtained from a tissue was not sufficient to ensure the reliability of flow cytometry data (One wild-type sample in Extended Data Fig. 2d; One control and one IL2 sample in Fig. 3b, 4f, Extended Data Fig. 7a, 9b, c, d; One mAb/IL2 sample in Fig. 4h). For survival plots shown in Figure 3a (NK1.1 depletion: One mouse on day two; Ly6G depletion: Three mice on day nine, five mice on day 16) and Figure 3b (CD4 depletion: One mouse on day nine; CD90.2 depletion: five mice on day seven) mice that needed to be sacrificed prematurely due to an acute intolerance towards the depletion antibody administration were censored from the analysis. In some experiments, single outliers identified by Grubb’s test were removed as indicated in the legends. Student’s unpaired t-test was used for the comparison of two means. If multiple groups were compared, one-way ANOVA was performed and if significant, multiple comparison was done by using Dunnet’s or Tukey’s post-hoc tests, except for tumor infiltration data, where comparison of multiple groups was done using Kruskal-Wallis test followed by Dunn’s test. If both time and treatment were compared, two-way ANOVA or a mixed model (if compared groups differed in size) was performed. If significant, multiple comparison was performed using Sidak’s post-hoc test. Significant differences in survival were determined using the log-rank test (Mantel-Cox) with Bonferroni correction for multiple comparisons. All statistical testes were two-tailed and performed with an alpha level of 0.05. GraphPad PRISM 8 was used for statistical analyses.

## Supplementary Figures

**Figure S1.**
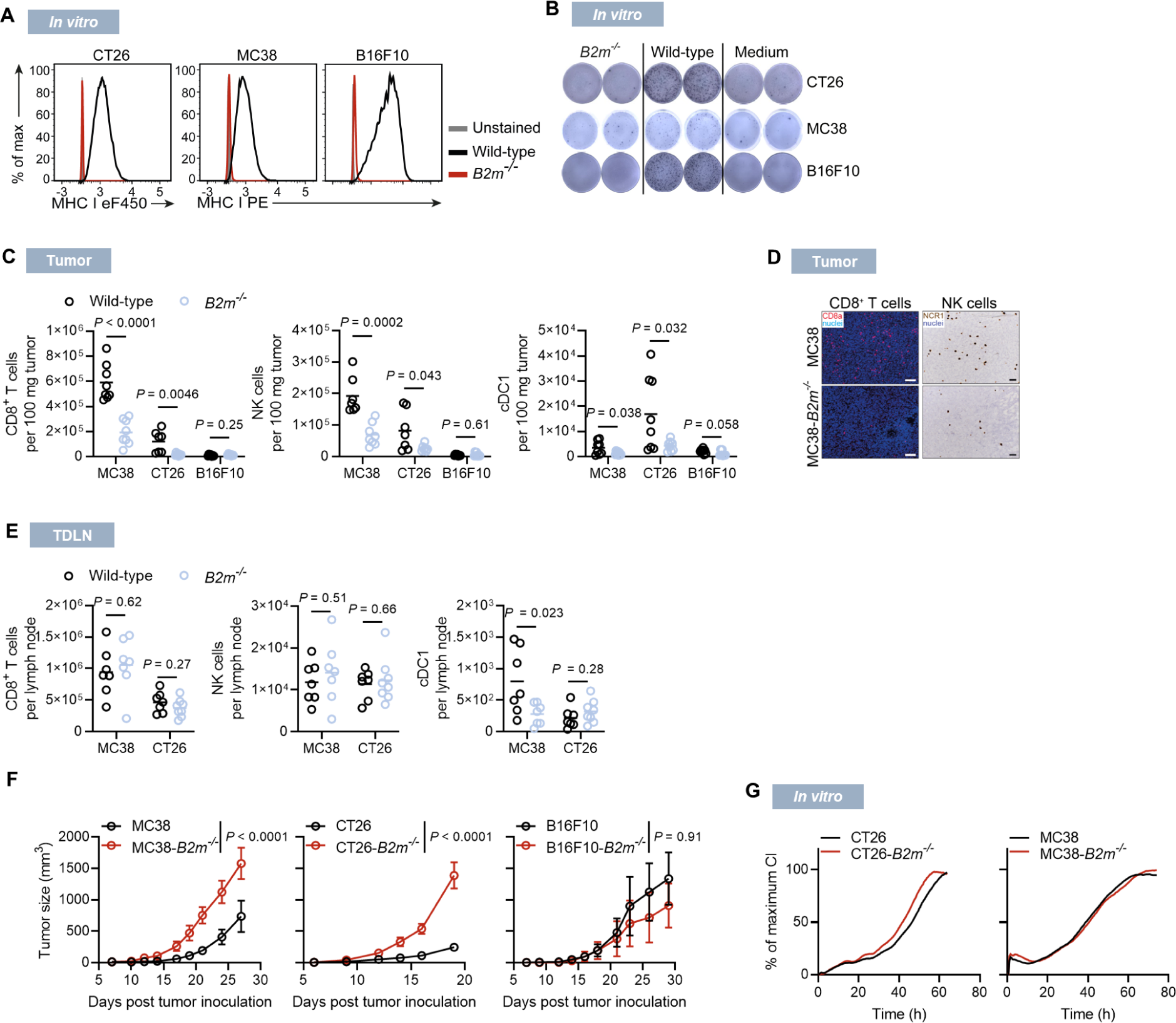
Characterization of MHC class I-deficient tumors. (**A**) MHC class I surface staining of wild-type and *B2m^-/-^* tumor cells. B16F10 cells were stimulated for 24 h with 25 ng/mL IFNγ prior to measurement. (**B**) IFNγ ELISpot of splenocytes from mice immunized with RNA-LPX encoding gp70 (CT26) or Ova257-264 (B16F10, MC38) co-incubated with wild-type and *B2m^-/-^* tumor cells as targets. B16F10 and MC38 cell lines were electroporated with Ova257-264-encoding mRNA. (**C** and **D**) Immune cell infiltration in CT26, MC38 or B16F10 wild-type and *B2m^-/-^*tumors that were grown in immunocompetent mice, determined by flow cytometry performed 20 d after inoculation (C) or by αCD8 staining and *Ncr1* ISH of MC38 wild-type and *B2m^-/-^* tumors 19 d after inoculation (D). Each row represents an individual tumor. Scale bar = 100 µm (IF) or 50 µm (*Ncr1* ISH). (**E**) Immune cell composition of TDLNs from mice bearing wild-type and *B2m^-/-^*tumors. (**F** and **G**) Growth of wild-type and *B2m^-/-^*cells *in vivo* (F) or in cell culture recorded by the xCELLigence system; CI, cell index (G). Single outliers were removed by Grubb’s test (C and E). n=8 (C). Representative examples (D). n=7-8 (E). n=5 (F). n=3 technical replicates (G).

**Figure S2.**
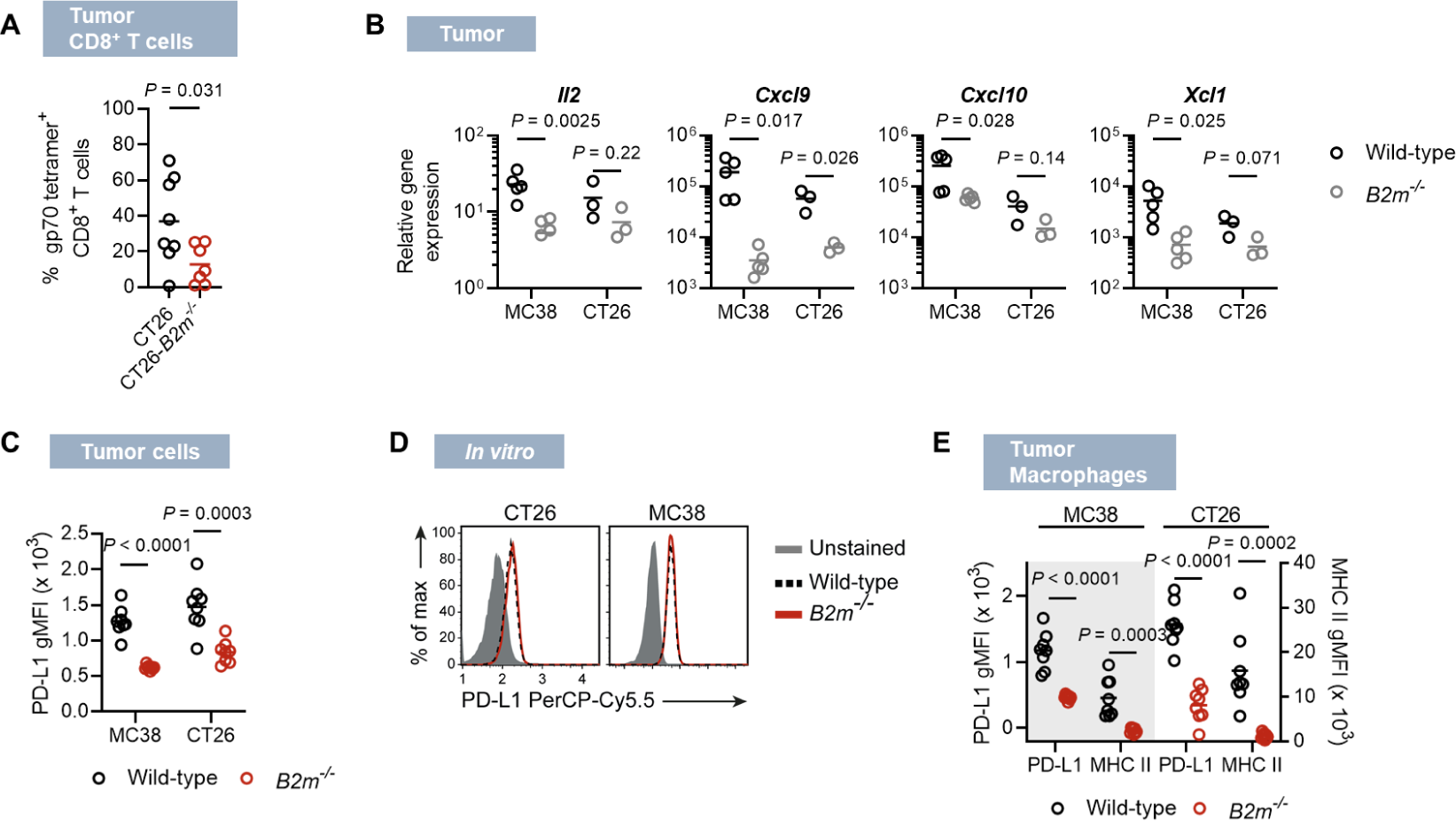
Inflammatory marker expression in MHC class I-deficient tumors. (**A**) Fraction of gp70-specific CD8^+^ T cells infiltrating CT26 and CT26-*B2m^-/-^* tumors 20 days after inoculation. (**B**) Gene expression pattern determined in wild-type or *B2m^-/-^* MC38 and CT26 whole-tumor tissues by qRT-PCR 19 d after inoculation. (**C** and **D**) PD-L1 expression in CT26 or MC38 wild-type compared to *B2m^-/-^* tumor cells *in vivo* at day 20 after inoculation (C) or after *in vitro* culture (D). (**E**) PD-L1 and MHC class II expression in macrophages infiltrating wild-type compared to *B2m^-/-^* tumors at day 20 after inoculation. n=7-8 (A). n=3 (CT26) and n=5 (MC38) (B). n=8 (C and E). Representative examples (D).

**Figure S3.**
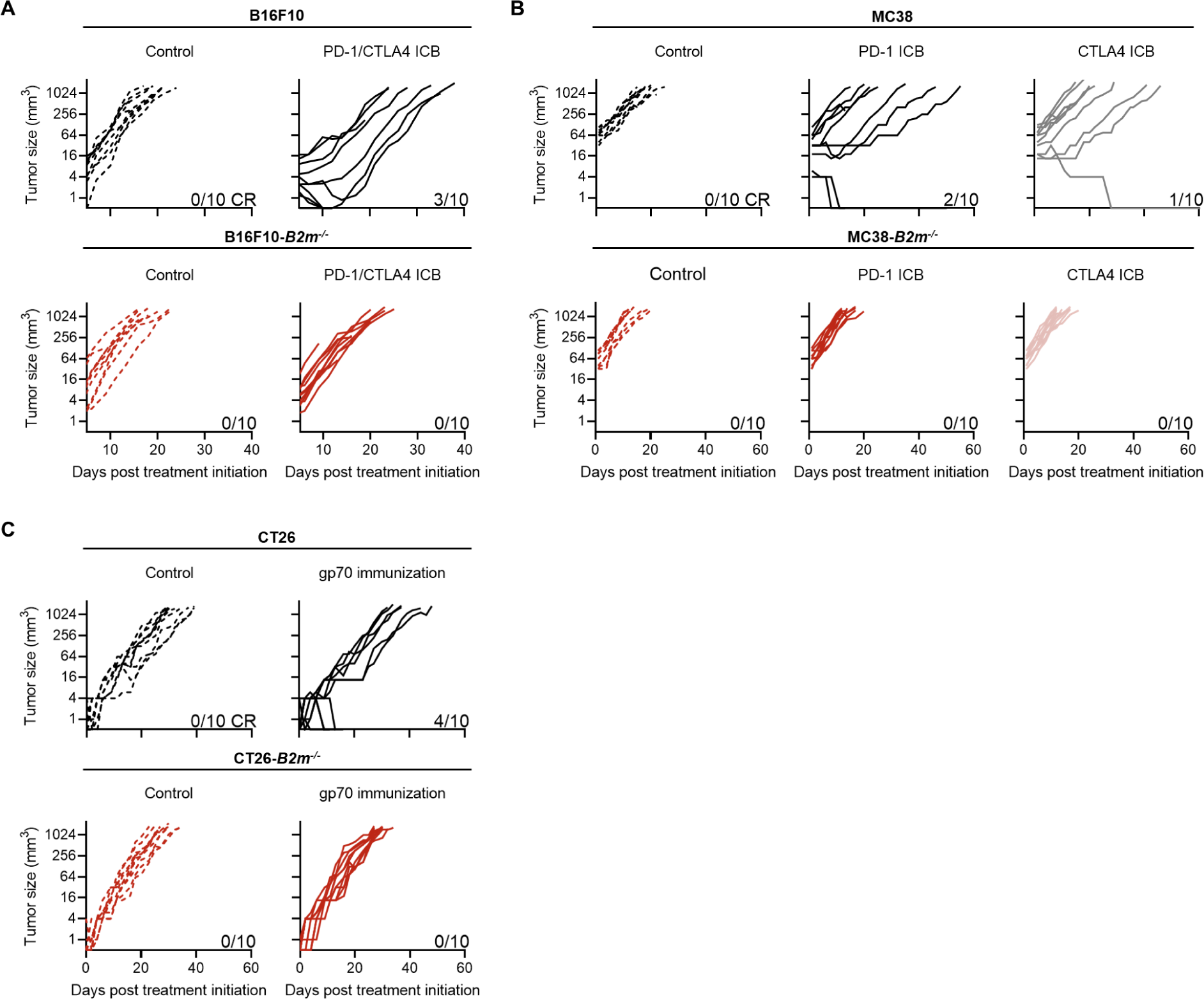
Treatment effect of T-cell based immunotherapies on MHC class I-deficient tumors. (**A** to **C**) Tumor growth curves of individual mice bearing wild-type or *B2m^-/-^* tumors treated with PD-1 and/or CTLA4 ICB or isotype controls (A and B) or with a gp70 mRNA-LPX vaccine or empty-vector mRNA-LPX as control (C). n=10 (A to C).

**Figure S4.**
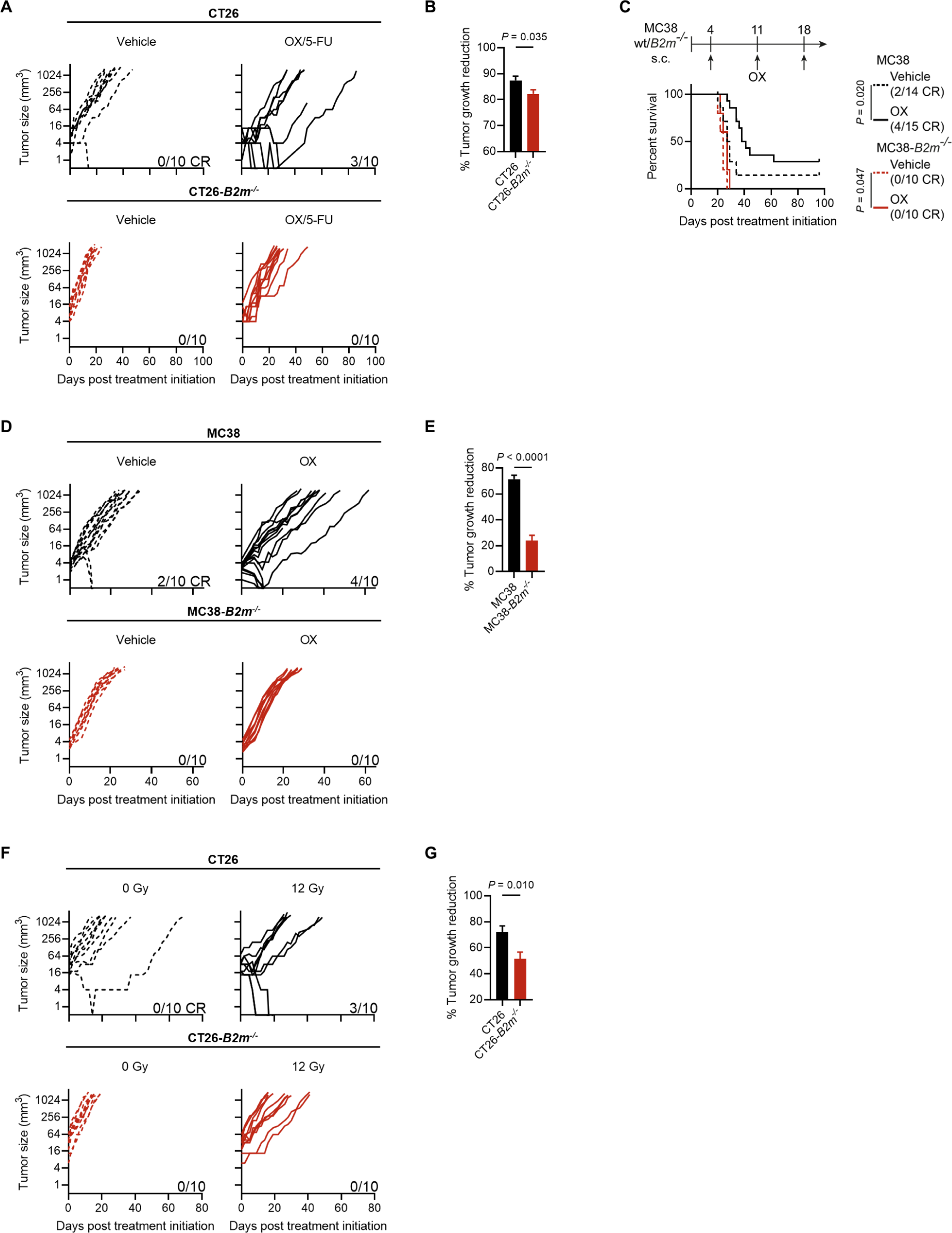
Treatment effect of chemo-and radiotherapy on MHC class I-deficient tumors. (**A** to **G**) Individual tumor growth curves (A, D and F), tumor growth reduction (B, E and G) and survival (C) of mice bearing wild-type and *B2m^-/-^* tumors. Mice were treated with Oxaliplatin/5- fluorouracil (OX/5-FU) doublet chemotherapy or vehicle as control (A and B), with OX or vehicle as control (C to E) or with Local radiotherapy (LRT) at a dose of 12 Gy or 0 Gy as control (F and G). Per cent growth reduction was calculated by dividing the average tumor volume of treated tumors through the average volume of control tumors and multiplying the resulting value with 100%. Error propagation was used to determine the s.e.m.. n=10 (A, B, F and G). n=10-15 (C to E).

**Figure S5.**
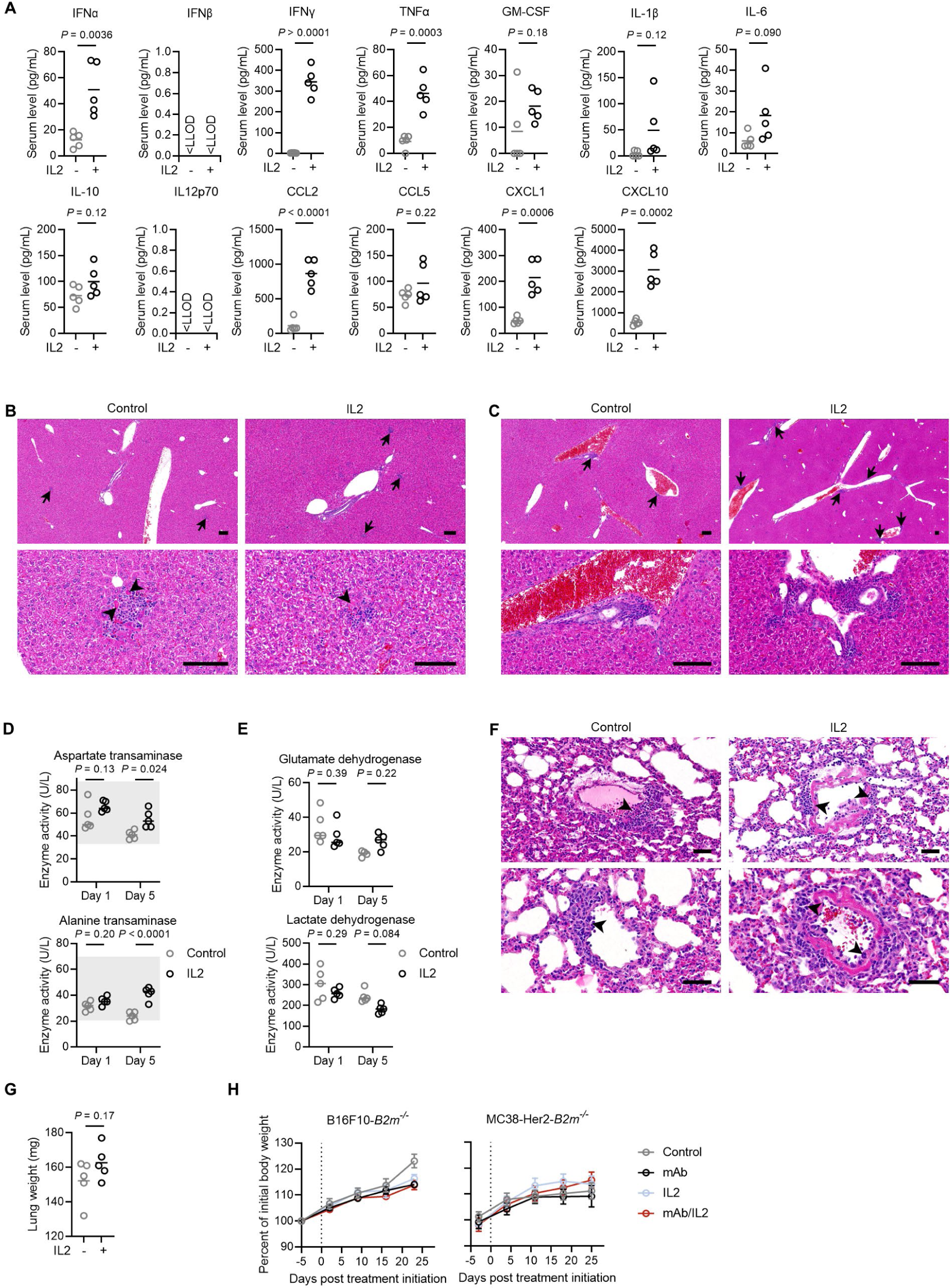
Toxicology of IL2. (**A-G**) Mice were treated once with intravenous Albumin-fused IL2 mRNA-lipopolyplex (short IL2) or with Albumin-encoding mRNA lipopolyplex as control. Cytokine induction in peripheral blood (A) was analyzed at day 1 after treatment. H&E-stained liver sections taken 5 days after treatment represent lobular (B) and perivascular and periportal (C) lesions, indicated by arrows, with arrowheads indicating single necrotic events. Scale bars represent 100 µm. Serum activity of transaminases (D) was analyzed 1 and 5 days after treatment. Reference ranges (shaded areas) of transaminases for naïve mice are 33.1-87.4 U/mL for aspartate transferase and 20.5-69.9 U/mL for alanine transaminase. Serum activity of dehydrogenases (E) was analyzed 1 and 5 days after treatment. H&E-stained lung sections (F) taken 5 days after treatment represent perivascular lesions, with arrowheads indicating immune cell infiltrates. Scale bars represent 100 µm. Lung weights (G) were analyzed at day 5 after treatment. (**H**) Body weight change of tumor-bearing mice treated as described in Fig. S6. n=5 (A, D, E and G). Representative images from an experiment with n=5 (B, C and F). n=10 (H).

**Figure S6.**
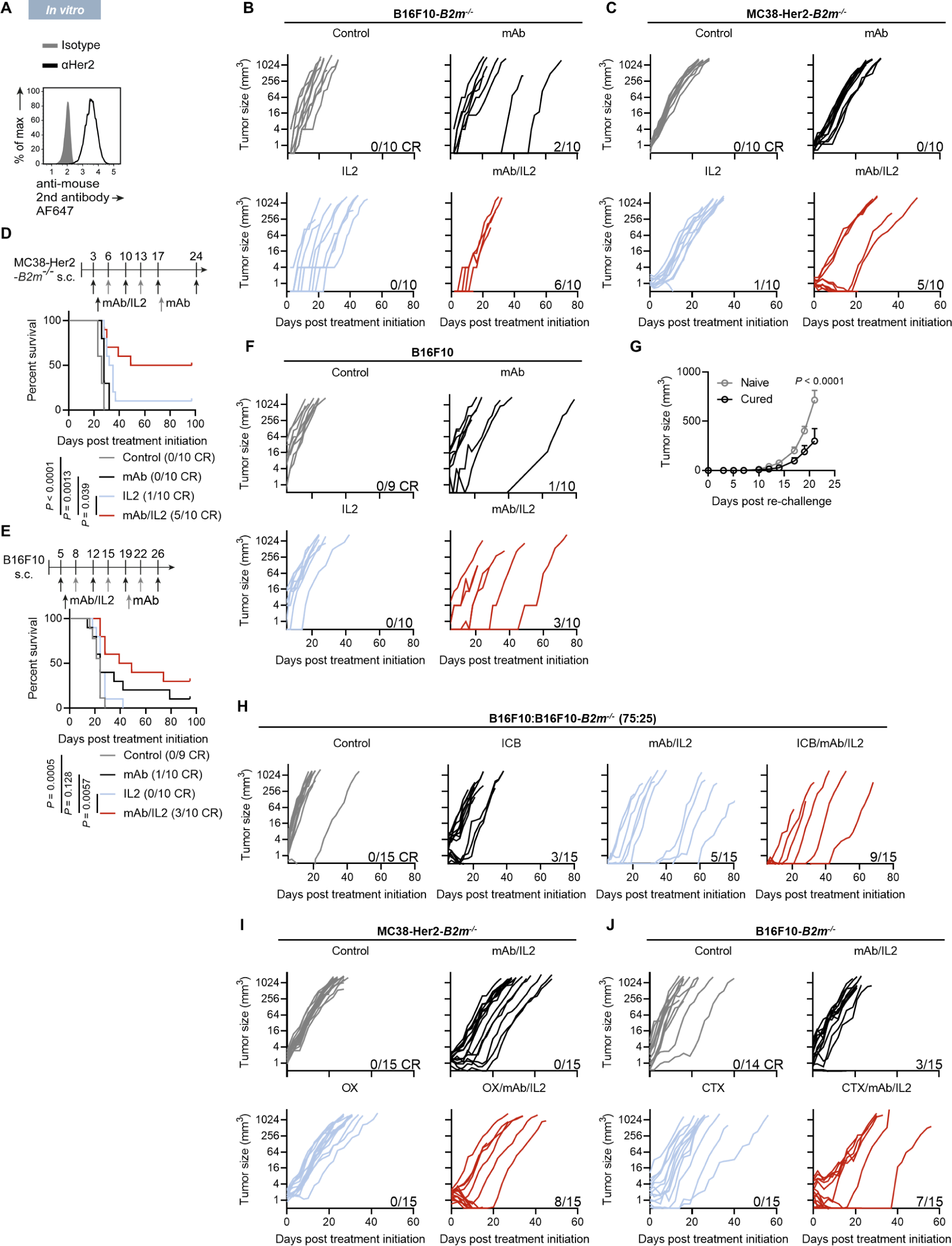
Treatment effect of mAb/IL2 on MHC class I-deficient tumors. (**A**) Her2 surface staining of MC38-Her2-*B2m^-/-^* cells. (**B** to **F**) Tumor growth curves of individual mice (B, C and F) and survival (D and E) for B16F10-*B2m^-/-^* tumors treated with αTrp1 mAb and intravenous IL2 mRNA-lipopolyplex (short IL2) (B), for MC38-Her2-*B2m^-/-^*tumors treated with αHer2 mAb/IL2 (C and D) and for B16F10 tumors treated with αTrp1 mAb/IL2 (E and F) when tumors were palpable. (**G**) Re-challenge of mice that rejected B16F10-*B2m^-/-^* tumors in response to mAb/IL2 treatment with B16F10-*B2m^-/-^* cells 72 days after initial tumor inoculation (53 days after treatment was concluded). Naïve mice served as controls. (**H**) Individual growth curves of heterogeneous B16F10 wild-type/*B2m^-/-^* tumors treated with αTrp1 mAb/IL2, PD-1 and CTLA4 combination ICB, or the quadruple combination. (**I**) Individual growth curves of MC38-Her2-*B2m^-/-^* tumors treated with αHer2 mAb/IL2, OX, or the triple combination. (**J**) Individual growth curves of B16F10-*B2m^-/-^* tumors treated with αTrp1 mAb/IL2, CTX, or the triple combination. Albumin-encoding mRNA and isotype mAbs served as controls across all experiments. Representative staining (A). n=10 (B to D). n=9-10 (E and F). n=6 (G). n=15 (H and I). n=14-15 (J).

**Figure S7.**
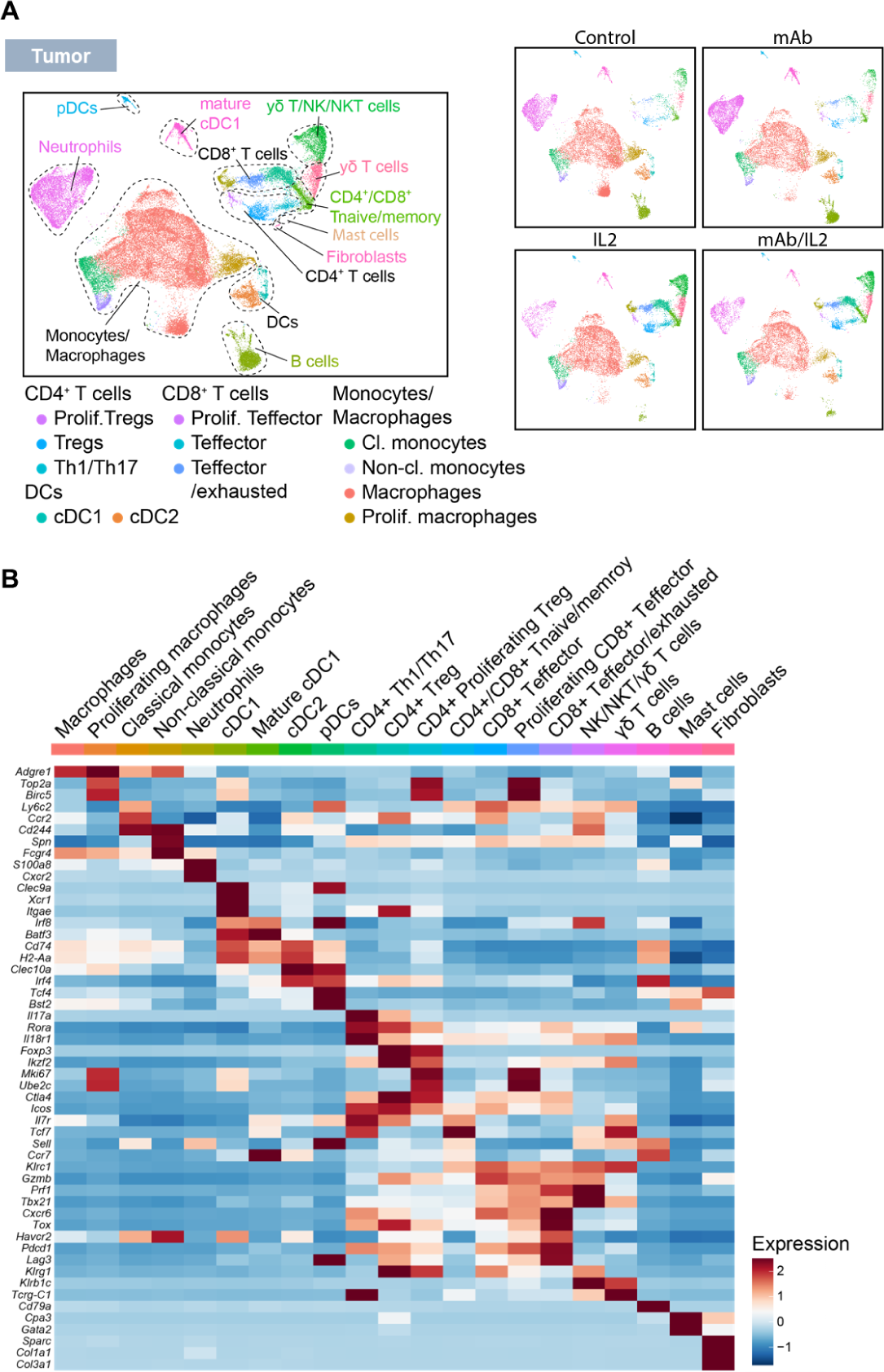
Immune cell subtypes infiltrating mAb/IL2-treated B16F10-*B2m^-/-^* tumors. (**A** and **B**) Mice were treated with αTrp1 mAb and intravenous IL2 mRNA-lipopolyplex (short IL2), either of the single compounds, or isotype control and albumin-encoding mRNA as controls at day 9 after inoculation. Tumor-infiltrating CD45^+^ cells were isolated 20 d after inoculation and subjected to scRNA-Seq. Merged (left) and treatment-specific (right) two-dimensional UMAP projection of a total of 52,794 analyzed cells (A) and expression of cell-type specific canonical markers across the different clusters (B). n=3 pooled per group.

**Figure S8.**
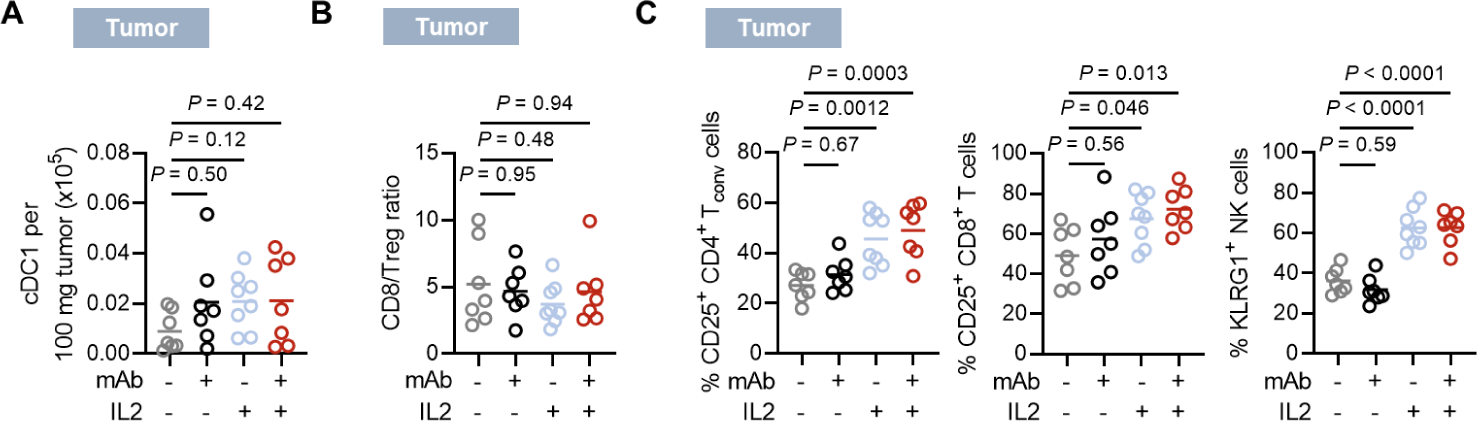
Infiltration and activation of immune cells in B16F10-*B2m^-/-^* tumors upon mAb/IL2 treatment. (**A** to **C**) Mice were treated with αTrp1 mAb and intravenous IL2 mRNA- lipopolyplex (short IL2), either of the single compounds, or isotype control and albumin-encoding mRNA as controls at day 9 after inoculation. Tumor-infiltrating leukocytes were analyzed 20 d after inoculation for cDC1 infiltration (A), CD8/Treg ratio (B) and expression of activation markers in CD4^+^ T cells, CD8^+^ T cells and NK cells (C). n=7 (mAb/IL2), n=7 (mAb), n=8 (IL2), n=7 (control) (A to C).

**Figure S9.**
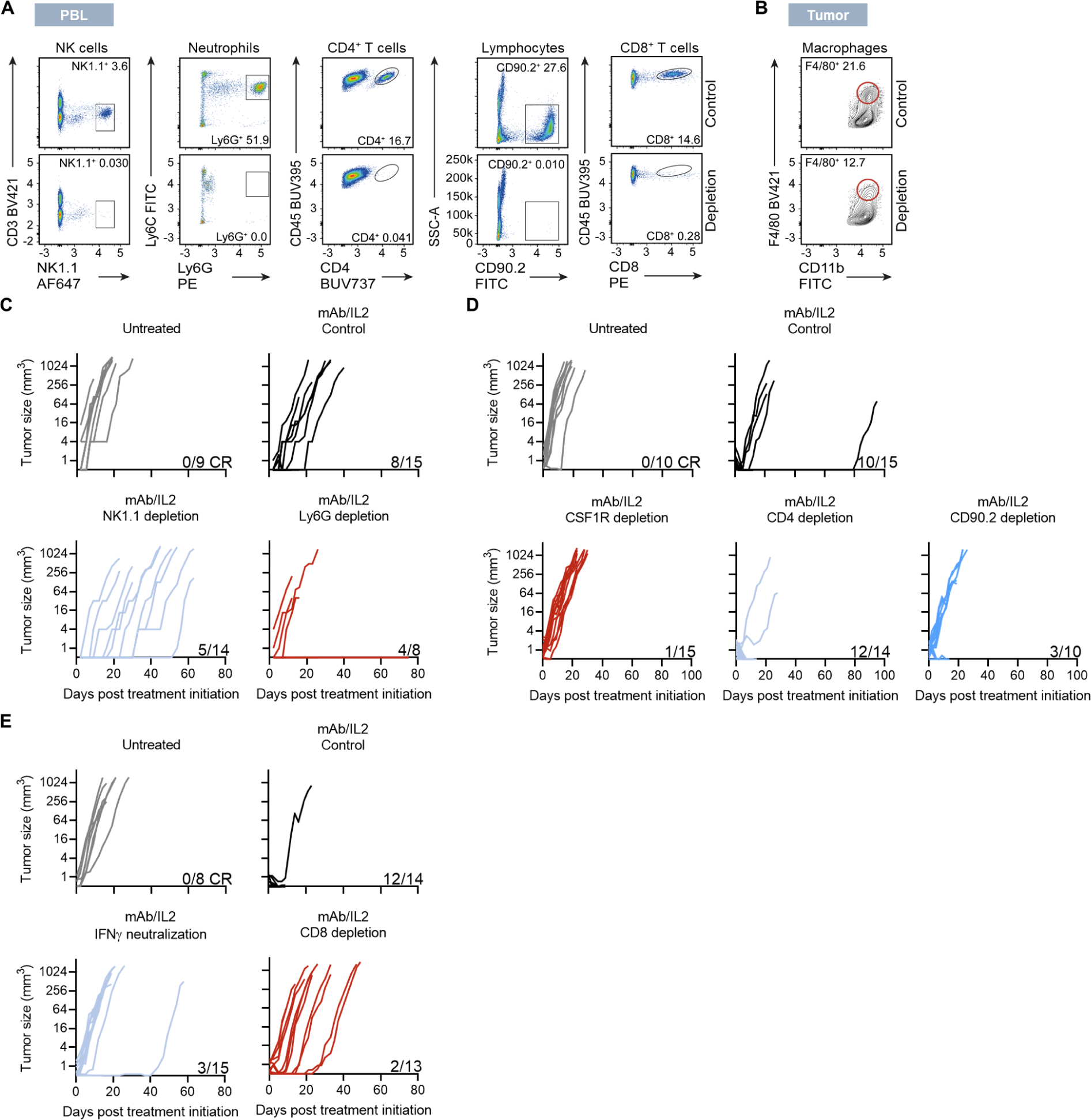
Identification of effector cells essential for anti-tumor activity of mAb/IL2. (**A** to **E**) Mice bearing B16F10-B2m^-/-^ tumors were treated with αTrp1 mAb and intravenous IL2 mRNA-lipopolyplex (short IL2), either of the single compounds, or isotype control and albumin-encoding mRNA as controls at day 9 after inoculation. Immune cell depletion in mice injected with depleting/neutralizing mAbs was analyzed by flow cytometry on the day of mAb/IL2 treatment initiation. Depletion of indicated cells was confirmed in blood (A) or tumor (B). Tumor growth curves of individual mAb/IL2-treated mice injected with mAbs against NK1.1 and Ly6G (C), CSF1R, CD4 and CD90.2 (D), or CD8 and IFNγ (E). Representative examples (A and B). n=9 to 15 (C to E). Representative examples (A and B). n=9-15 (C). n=10-15 (D). n=8-15 (E).

**Figure S10.**
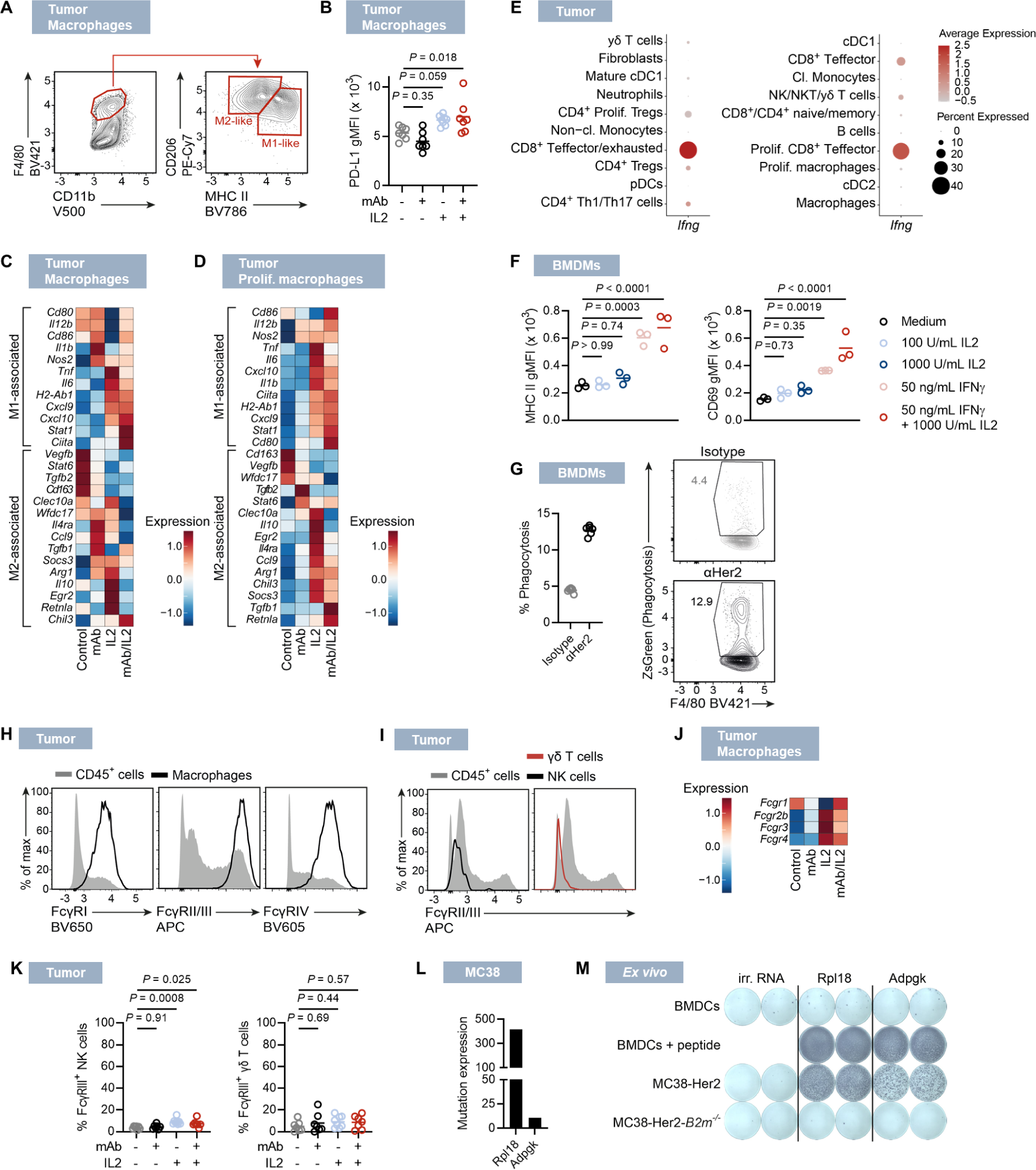
Characterization of effector cells infiltrating mAb/IL2-treated B16F10-*B2m^-/-^* tumors. (**A** to **E**) Mice bearing B16F10-*B2m^-/-^* tumors were treated with αTrp1 mAb and intravenous IL2 mRNA-lipopolyplex (short IL2), either of the single compounds, or isotype control and albumin-encoding mRNA as controls at day 9 after inoculation. Macrophages infiltrating the tumors were analyzed with regards to M1/M2 polarization discriminated by CD206 and MHC class II expression (A), PD-L1 expression (B) and expression of selected M1- or M2-associated genes in cell clusters representing macrophages (C) and proliferating macrophages (D) as determined by scRNA-Seq. Dot plot showing expression of *Ifng* in leukocytes from B16F10-*B2m^-/-^* tumors treated with αTrp1 mAb/IL2 (E). (**F**) BMDM were stimulated for 24 h with 50 ng/mL IFNγ, 1000 U/mL IL2, a combination of both or cultured without additives as control and analyzed for expression of MHC II and CD69. (**G**) Quantification of ZsGreen incorporation of BMDM stimulated with IFNγ as described in (F) and subsequently co-cultured with ZsGreen mRNA-electroporated MC38- Her2-*B2m^-/-^* cells opsonized with αHer2 mAb or isotype control. (**H**) Expression of FcγRs in B16F10-*B2m^-/-^* tumor-infiltrating macrophages 20 d after inoculation. (**I**) Expression of FcγRs in B16F10-*B2m^-/-^* tumor-infiltrating NK cells and γδ T cells 20 d after inoculation. (**J** and **K**) Tumors were treated as described in (A). The expression of genes encoding FcγRs in infiltrating macrophages was analyzed by scRNA-Seq 20 d after inoculation (J) and FcγR expression by NK cells and γδT cells was analyzed by flow cytometry (K). (**L**) Expression of Rpl18 and Adpgk neoepitopes in cultured MC38 cells. (**M**) CD8^+^ T cells from mice immunized with the indicated RNA-LPX were subjected to IFNγ ELISpot using BMDC and mutant antigen peptides or tumor cells. Representative examples (B, H and I). n=3 pooled per group (E-D and J). n=7-8 (B and K). n=3 (F). n=6 technical replicates (G).

## Notes

### Competing Interest Statement

J.D. Beck, M. Vormehr, L. Allnoch, M. Diken and S. Kreiter are employees and OE. Tuereci and U. Sahin are cofounders and management board members at BioNTech SE (Mainz, Germany). J.D. Beck, M. Vormehr, L. Allnoch, M. Diken, S. Kreiter, OE. Tuereci and U. Sahin hold securities from BioNTech SE. J.D. Beck, M. Vormehr, M. Diken, S. Kreiter and U. Sahin are inventors on pa-tents or patent applications related to this study.

### Summary of Updates

Section headings. Fig. 4 became overloaded and was split into Fig. 4 and Fig. 5. Experimental data and a more detailed description of the IL2 mRNA biodistribution (page 5). The language was revised throughout the manuscript to better explain the mode of action, in particular the role of IL-2 mRNA in driving the tumor-specific CD8+ T-cell response. A re-challenge study showing reduced growth of B16F10-B2m-/- cells in mice cured from B16F10-B2m-/- tumors was performed and included (Fig. S6G). A paragraph related to IL-2 toxicity (page 6) including data from additional experiments comprising representative histopathological images of lungs and livers, clinical chemistry, lung weights, and cytokine induction was provided (Fig. S5). Dendritic cell infiltration (Fig. S8A) and CD8/Treg ratio (Fig. S8B) in mAb/IL2 treated tumors were investigated and included. Enhanced cross-presentation of tumor-derived antigen by macrophages upon IL2 treatment (Fig. 5A) was studied and included. Investigation of Fcγ receptor expression in tumor-infiltrating NK cells and γδ T cells (Fig. S10, I and K) was provided.

